# Patient-Derived Organoids Recapitulate Intrinsic Immune Landscapes and Progenitor Populations of Glioblastoma

**DOI:** 10.1101/2021.10.06.463228

**Authors:** Fumihiro Watanabe, Ethan W. Hollingsworth, Jenna M. Bartley, Lauren Wisehart, Rahil Desai, Annalisa M. Hartlaub, Mark E. Hester, Paula Schiapparelli, Alfredo Quiñones-Hinojosa, Jaime Imitola

## Abstract

Glioblastoma stem cells (GSCs) are highly self-renewing, resistant to therapy, and are able to form lethal tumors^1, 2^. Tumor organoids have been developed to study tumor evolution^1–4^, and while GSCs can form organoids for glioblastoma multiforme, our understanding of their intrinsic immune, metabolic, genetic, and molecular programs is limited. To address this, we deeply characterized GSC-derived GBM organoids using a modified protocol (GBMOsm) from several patient-derived GSCs and found they develop into complex 3D tissues with unique self-organization, cancerous metabolic states, and burdensome genetic landscapes. We discovered that GBMOsc recapitulate the presence of two important cell populations thought to drive GBM progression, SATB2^+^ and HOPX^+^ progenitors. Despite being devoid of immune cells, transcriptomic analysis across GBMOsc revealed an immune-like molecular program, enriched in cytokine, antigen presentation and processing, T-cell receptor inhibitors, and interferon genes. We determined that SATB2^+^ and HOPX^+^ populations contribute to this immune and interferon landscape in GBM in vivo and GBMOsm. Our work deepens our understanding of the intrinsic molecular and cellular architecture of GSC-derived GBMO and defines a novel GBMOsc intrinsic immune-like program.

## Introduction

Glioblastoma multiforme (GBM) remain one of the most lethal cancers worldwide^5^. One prevailing model of GBM posits that tumors originate and recur from glioblastoma stem cells (GSCs)^5–7^. Like other stem cells, GSCs have increased self-renewal capacity and exhibit a progenitor-like state, but differ in their pathological properties, including increased survival, aneuploidy, oncogenesis, and resistance to therapy. Beyond the cell-autonomous component of GSCs and maturing cancer cells, GBM evolve by forming a tumor microenvironment (TME). That is, a collection of non-tumoral cells, including vascular, immune, and mesenchymal cells, that permit angiogenesis, local immunosuppression (thereby facilitating tumor cell proliferation), and migration and invasion into normal tissue, which altogether leads to explosive tumor growth. This TME crosstalk shapes the intrinsic properties of the developing tumor itself. One important example, for instance, is that GBM cells regulate infiltrating cell types according to their own mutational landscape^8^, as well as their own glioma-intrinsic gene expression programs^9^. Thus, how exactly an initial cluster of GBM cells can evolve and co-opt the TME to favor tumor promotion remains a fundamental unanswered question with important clinical and therapeutic implications.

Preclinical models have consisted mainly of genetically engineered mouse models or human tumor-derived cell lines propagated in either two-dimensional (2D) cultures or as patient-derived xenografts (PDXs) in mice^10, 11^. There has been a growing interest in glioblastoma organoids^4, 12–16^. Organoids are defined as 3D *in vitro* tissue-like constructs derived from isolated stem cells which mimic their corresponding *in vivo* organ. Among the several models of GBM organoids ^4, 12–16^ is one derived from intact microscopic pieces of tissue from surgically resected tumors termed (GBO). This model offers the advantage of closely replicating its native tumor, since the tissue has not undergone major alterations other than being placed in defined cultures, and thus could be very useful for chemical screenings of potential therapeutic compounds. However, this model is less suitable for studying the initial GBM intrinsic program and early self-organization of GBM since the tumor is already organized *in vivo* and retains its complete TME, with infiltrating immune and vascular cells^14^. Therefore, it is challenging to experimentally distinguish the intrinsic human GBM from the stroma laden with diverse cells. Moreover, by the time GBM are usually detected, tumor and microenvironment interactions are long established. An alternative GBM organoid model is one derived from GSCs isolated from a patient’s resected GBM surgical tissue, which are used to generate GBM organoids, recapitulating aspects of GBM tissue and organization^15^. However, it relies on the addition of exogenous growth factors during organoid formation and differentiation, which may favor unwanted proliferation of GSCs and clonal selection rather than inducing the initial events of 3D tissue-like formation seen in regular iPSCs and NSCs-derived organoids and making comparison difficult, since these organoids don’t require growth factors during the organoid formation stage.

In this study, applying an improved protocol, we performed a comprehensive, integrative characterization of GBM organoids generated from several patient-derived GSCs, we termed GBMOsm (GSC-derived <u>GBM O</u>rganoid derived from GBM <u>S</u>tem Cells with <u>M</u>odified Protocol), Henceforth, GBMOs. We specifically examined GBMO microanatomy, progenitor diversity, mutational and transcriptomic landscapes, and their intrinsic gene expression programs, especially as they relate to GBM *in vivo*. Our work advances our understanding of the intrinsic cellular and molecular features of GSC-derived GBMO, including the identification of novel progenitor populations and immune gene signatures.

## Results

### Patient-derived GSCs self-organize into 3D tumor tissue-like structures

GBM have unique spatial and temporal characteristics, with tissue-like domains and cellular diversity^15^. GSCs are also thought to underlie glioma origin, initiation, evolution, and therapy resistance. Studying GSCs is usually done by injecting these cells into mice brains, as a xenotransplant into a immunosuppressed host. This approach is less suitable to dissect autologous neuroimmune interactions^17^ seen in neuroinflammation^5, 18^ and in brain cancer^7^. Previously, we showed the intrinsic ability of normal neurospheres to form through aggregation, a complex tissue-like construct, resulting in self-organization, and progenitor diversity that we called NEDAS^19^. Furthermore, we have shown that neurospheres have an intrinsic immune program, which express T cell co-stimulatory molecules and chemokine receptors that are functional during neuroinflammation in multiple sclerosis models^20–22^. Thus, we hypothesized that GSCs spheres like NSCs spheres may share developmental potential characteristics. A previous report showed GSCs are able to form organoids, but the study of their intrinsic properties including their neuroimmune phenotype is unknown^15^.

To expand on these efforts, we cultured GSC-derived tumorspheres from patients who underwent GBM resection and then generated GBMOs, that are proven to be GSCs *in vivo* and *in vitro* in prior work.^23–26^ (**Supplementary Figure 1a-c, Supplementary Table 1**). Prior GSC-derived organoids were maintained at all times under conditions of EGF/FGF. Typically organoids are maintained in defined medium without added growth factors from organoid generation, we favored this method to allow comparison with control organoids. We embedded GSCs in Matrigel and generated GBMOs with a 94% success rate (n=314 organoids from 6 GSC lines after 4 weeks in culture). **Supplementary Fig. 1f, 2a-b** The organoids reached sizes 20-fold larger than GSCs grown as “tumorspheres” (a cancer cell aggregate;). (**Supplementary Fig. 2a,b).** GSCs aggregated and integrated into a uniform tissue architecture. Pathology comparison of GBMOs and parental tumors showed proliferative and quiescent/necrotic compartments, in addition to the presence of pseudopalisading cells in selected tumor-organoid pairs (**Supplementary Fig. 1d,e**). For instance, GBMO-965 remained sparsely cellular with syncytia-like formations while GBMO-1201 retained its multicellularity with fibrils and occasional highly chromatic nuclei. These findings are consistent with recent work indicating that GBM organoids derived from intact tissue (GBOs) reproduce features of their parental tumors^15^. These findings demonstrated that GSCs can form 3D organoids without added growth factors allowing us to perform an in depth-phenotypic characterization and comparisons using similar cultures protocols for iPSCs (iPSCO) and organoids derived from NSCs (NSCO).

### Patient-derived GBMO host distinct microenvironments and progenitor diversity mimicking GBM biology

Next, we compared GBMO self-organization to non-tumoral, age-matched organoids derived from healthy neurospheres from H9-derived human neural stem cells (NSCO)^19^ (**Fig.1a)**. We also used cerebral organoids derived from human iPSCs (iPSCO), which we and others have shown recently to model the stereotypical architecture of the developing brain with self-organized progenitor niches^27, 28^ and cortical layering. GBMO, NSCO, and iPSCO, all maintained in Matrigel and growth factor-free culture conditions, uniformly expressed the human NSC marker, vimentin (VIM)^29^. These organoids were composed of segregated areas of proliferation, marked by Ki-67 (Ki-67^+^), apoptosis by activated-caspase-3 (A-Cas^+^), and quiescent/stressed cells by activating transcription factor 4 (ATF4^+^), albeit to a lesser extent in NSCO (**Fig. 1b-f and Supplementary Fig. 3a, b**). By contrast, GSCs cultured as tumorspheres in EGF/FGF lacked any such organization (**Supplementary Fig. 2a, b**). We quantified Ki-67^+^ and A-Cas^+^ layer thickness among GBMO and found GBMO-30 contained marked overlap while the others exhibited a clear delineation of proliferating cells from the outside surface and apoptotic and quiescent cells from the inner core (**Fig. 1c-e**). As organoids expand, their inner core experiences a reduction in oxygenation due to diffusion limits^15^. To determine hypoxic gradients in GBMOs, we stained for the BCL-2 interacting protein-3 (BNIP3), a protein downstream of hypoxia-inducible factor 1α (HIF-1α) and a marker for hypoxia^30^. We found increased cytoplasmic and nuclear BNIP3 expression inside GBMO compared to their surfaces, as shown by quantitative confocal intensity profile measurements (**Fig. 1b and Supplementary Fig. 3c, d, e, f)**. Collectively, these data indicate that all examined GBMO hosted microenvironments for apoptosis, proliferation, hypoxia, and cellular stress as seen in GBM *in vivo*.

**Figure 1.**
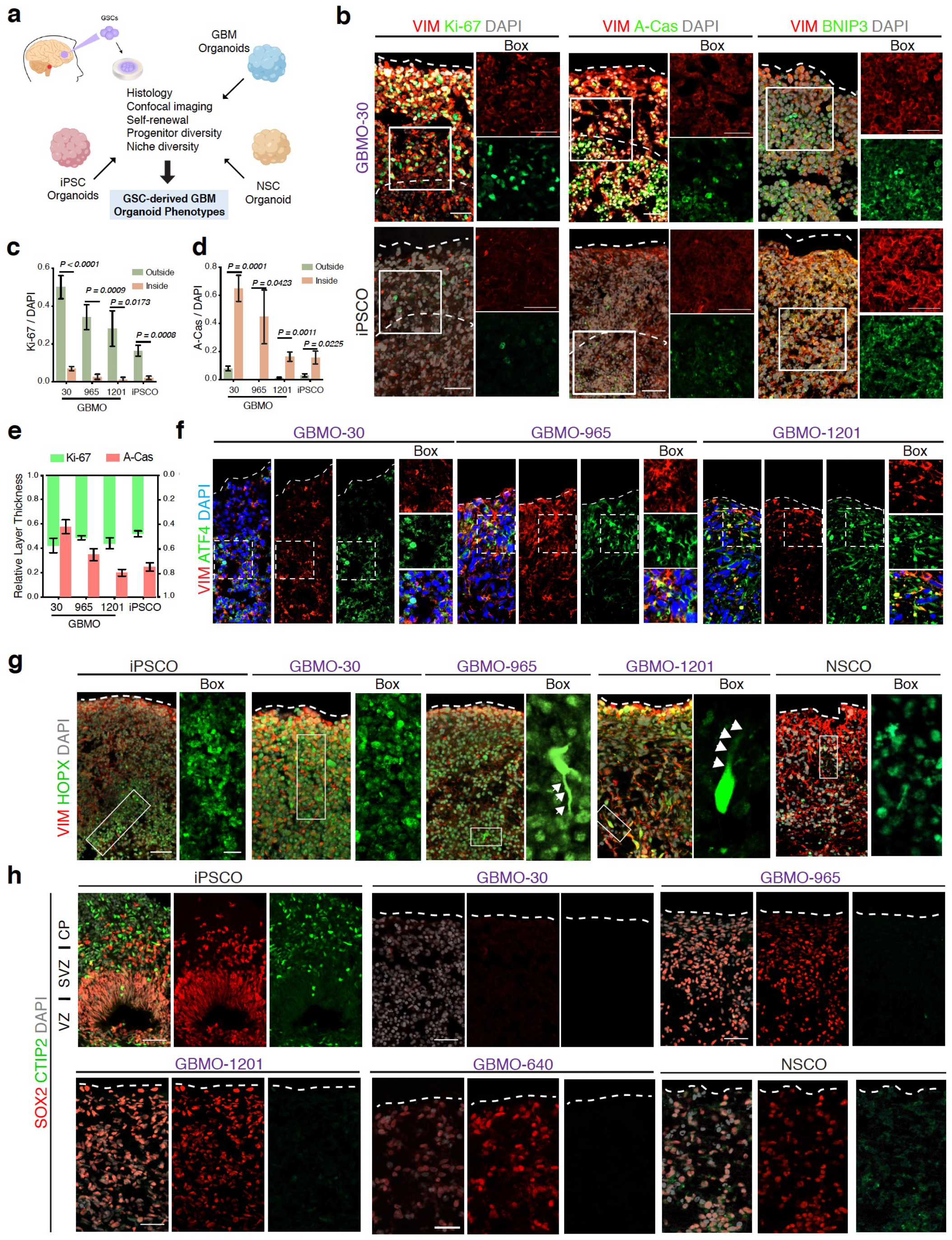
Deep characterization of GBMO defines layering organization, patient-specific differences, and partial co-optation of a developmental program. **(a)** Approach for studying patient-derived glioblastoma in the GBM organoid model. **(b)** Representative Immunohistochemistry for vimentin (VIM), proliferative (Ki-67), apoptotic (active-caspase-3 (A-Cas)), and hypoxia (BNIP3) markers in GBMO-30 and induced pluripotent stem cell organoid (iPSCO). Scale bar, 50 μm. (c-e) Quantification of Ki-67-positive **(c)** and A-Cas-positive **(d)** cells, and relative thickness of proliferative and apoptotic areas **(e)** in iPSCO and patient-derived GBMOs. Bar graphs are presented as means ± SEM. Two-tailed unpaired Student’s t-test. (Ki-67: GBMO-30, *P* < 0.0001; GBMO-965, *P =* 0.0009; GBMO-1201, *P =* 0.0173; iPSCO, *P =* 0.0008. A-Cas, GBMO-30, *P* = 0.0001; GBMO-965, *P =* 0.0423; GBMO-1201, *P* = 0.0011; iPSCO, *P =* 0.0225.) **(f)** Immunofluorescent images for quiescent/stress marker ATF4 in GBMO-30, 965 and 1201. **(g)** Sample images of immunostaining for VIM, and HOPX markers of outer radial glia in iPSCO, GBMO-30, −965, −1201, and neural stem cell organoid (NSCO). Scale bar, 20 μm. **(h)** Immunostaining for the stem cell marker SOX2 and deep-layer neuron marker CTIP2 in iPSCO, NSCO, and GBMO lines. Scale bar, 50 μm.

To determine the functional basis for the expression of the radial glia marker VIM in GBMOs, we stained for the phosphorylated form (p-VIM), a marker for radial glia division. Notably, we found a population of p-VIM^+^ radial glia-like tumor cells undergoing mitosis in the proliferating layers of GBMO (**Supplementary Fig. 3g**). We confirmed their identity by staining for homeodomain-only protein homeobox (HOPX), a transcriptional regulator and marker for outer radial glia (oRG) in normal human neurodevelopment. Unlike their stereotypical localization to the outer subventricular zone (SVZ) in iPSCO, we observed multiple HOPX^+^ cells scattered throughout the outer edge of GBMOs, some with characteristic polarity and elongated processes (**Fig. 1g**). This oRG-like population in GBM has been recently described as invasive GBM cells with stem cell and migratory properties *in vivo*. Altogether, these data demonstrate that GBMO derived from GSCs are capable of replicating the pathological oRG-like cell population observed in GBM *in vivo* and GBM tissue *ex vivo*^14, 31, 32^.

The identification of these oRG-like tumor cells, together with the known ability of GSCs to co-opt developmental programs to direct tumorigenesis^33^, prompted us to examine the molecular diversity of progenitor markers in GBMO. We performed an analysis of well-defined neural progenitor markers in all GBMO and compared them with their expression levels reported in GBM *in vivo* (**Fig 2b**). In iPSCO, as expected, we observed well-defined SOX2^+^ ventricular zone (VZ) and SVZ-like areas adjacent to cortical plate-like areas containing CTIP2^+^ cells, a marker for early born cortical neurons^27^ (**Fig. 1h**). In contrast, we did not observe CTIP2^+^ cells in any GBMO, mirroring its low expression in GBM *in vivo* (**Fig. 2b**). Though unsurprisingly absent in GBM-30, which is characterized by the mesenchymal phenotype *in vivo*^24, 25, 34^, SOX2^+^ cells were variable among patient-derived lines and dispersed throughout the layers of GBMO (**Fig. 1h**). We stained for additional lineage markers, including PAX6; the intermediate cell marker TBR2; deep-layer neuron markers, SATB2 and TBR1; and Cajal-Retzius cell marker, REELIN (**Fig, 2a**). Unlike iPSCO, where cells were stereotypically located, these cell type markers showed disorganized expression scattered throughout GBMO. Marker expression intensity largely reflected what has been reported in GBM *in vivo*. For instance, SATB2 (**Fig. 2a-middle**), a bona-fide marker for GSCs that was recently found to be a driver for GBM growth^35^, was highly expressed in GBMO compared to NSCO, while TBR1 was expressed in all GBMOs (**Fig. 2a-upper**). Collectively, these results indicate that GSCs form organoids that not only contain proliferative, apoptotic, and quiescent niches that vary in their relative size, but also reactivate a progenitor program like GBM *in vivo*, including the oRG-like HOPX^+^ and SATB2^+^ cancer stem cell populations (**Fig. 2b,c**).

**Figure 2.**
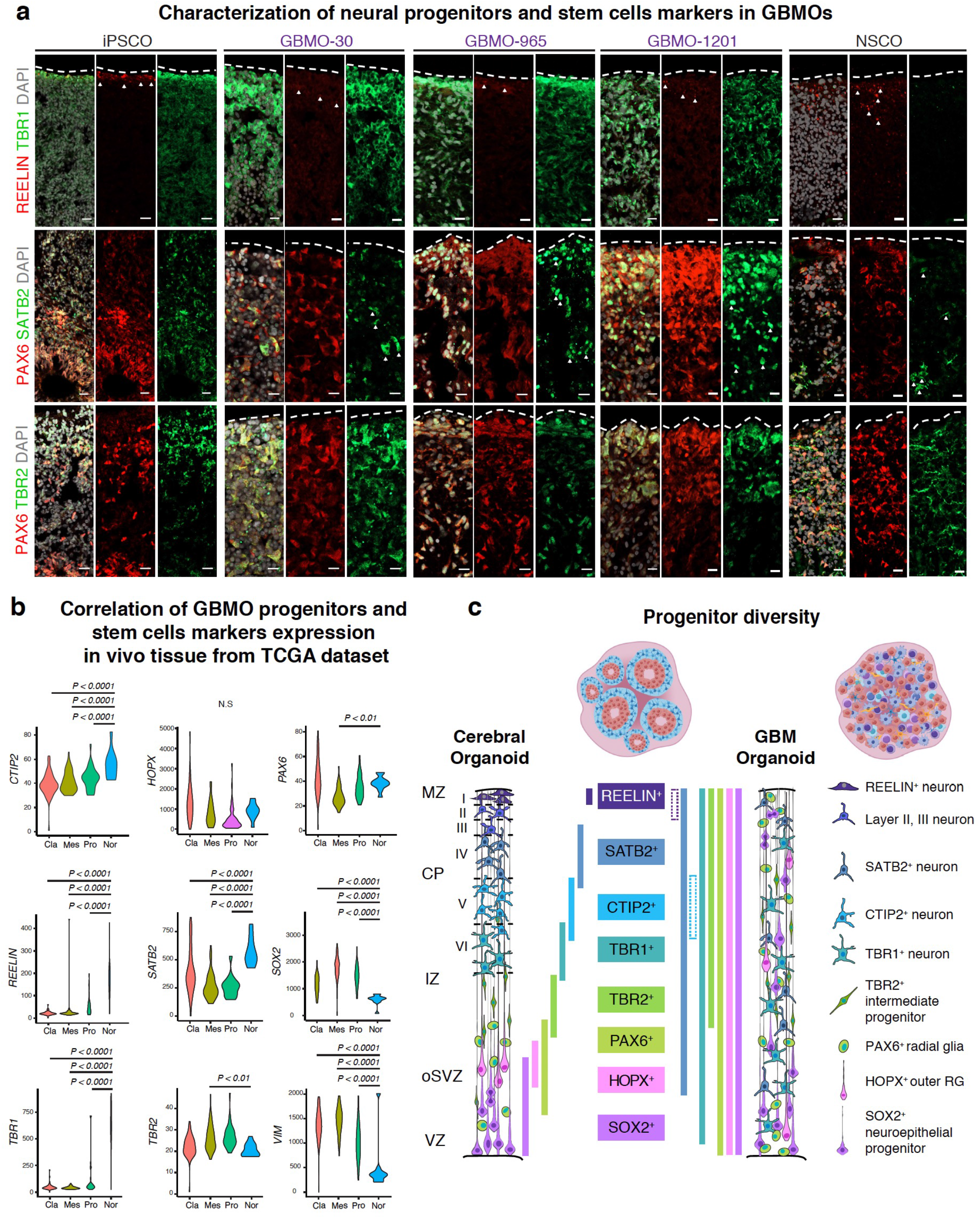
Characterization of markers for progenitor diversity in GBMO and their expression in vivo. **(a)** Representative confocal images for immunostaining of Cajal-Retzius cell (top: REELIN), deep-layer cortical neuron (top: TBR1, second: SATB2), intermediate progenitor (second: TBR2), and radial glia (second and third: PAX6) markers in iPSCO, hNSO, and each GBMO line. Scale bar, 20 μ**(b)** Violin plots of marker mRNA expression by molecular subtype using publicly available TCGA cohorts. Cla, Classical; Mes, Mesenchymal; Pro, Proneural; and Nor, Normal. One-way ANOVA followed by Dunnett’s correction, relative to Normal. (*CTIP2*: Cla, *P* < 0.0001; Mes, *P* < 0.0001; Pro, *P* < 0.0001. *PAX6*: Mes, *P* < 0.01. *REELIN*: Cla, *P* < 0.0001; Mes, *P* < 0.0001; Pro, *P* < 0.0001. *SATB2*: Cla, *P* < 0.0001; Mes, *P* < 0.0001; Pro, *P* < 0.0001. *SOX2*: Cla, *P* < 0.0001; Mes, *P* < 0.0001; Pro, *P* < 0.0001. *TBR1*: Cla, *P* < 0.0001; Mes, *P* < 0.0001; Pro, *P* < 0.0001. *TBR2*: Mes, *P* < 0.01. VIM: Cla, *P* < 0.0001; Mes, *P* < 0.0001; Pro, *P* < 0.0001.). N.S. = not significant, *P* > 0.05. **(c)** Schematic of tissue-like organization and progenitor diversity in iPSCO and GBMO, color bands represent the localization or absence of specific progenitor organized in layers in the normal organoids compared to glioblastoma organoids.

### Genomic architecture of GBMO reflect alterations found in GBM

Thus far, detailed genomic and mutational analyses of GSC-derived GBMO have not been performed. *In vivo*, GBM present with a high degree of aneuploidy, copy number changes, and somatic mutations^36^. To evaluate the genetic landscape of GBMO, we performed targeted massively parallel DNA sequencing (TMPDS) and large-scale chromosomal copy-number variation (CNV) analysis of all patient-derived GBMOs^37^ and NSCO^19^ that were cutured simultenously with identical culture conditions **(Fig.3a).** CNV analysis revealed a greater number of CNVs across all GBMOs, relative to NSCO. These included sizable amplifications in chromosome 7 and deletions in chromosome 10, which were shared among GBMO-1201, GBMO-965, and GBMO-640 and have been frequently reported in GBM (**Fig. 3b and Supplementary Table 4**). We next employed TMPDS to identify somatic mutations in GBMO and NSCO. From a panel of 407 cancer-related genes, we identified 32 somatic mutations in GBMO-30, nine in GBMO-965, seven in GBMO-1201, and five in GBMO-640 while NSCO harbored five non-damaging mutations (**Fig. 3c and Supplementary Table 5**). Of the genes mutated in GBMO, the majority (GBMO-30, 70%; GBMO-965, 89%; GBMO-1201, 71%; GBMO-640, 60%) are mutated in native GBM *in vivo*^38^. Moreover, whereas four of the five mutated genes in NSCO are tolerated missense (SIFT score <0.05) (**Fig. 3d**), most GBMO mutations are predicted to be damaging, frame-shifting, or protein-truncating (**Fig. 3c-e**). GBMO-30, in particular, was composed of the most damaging frame-shift and nonsense mutations (**Fig. 3d,f**).This mutational spectrum, consisting of nonsynonymous mutations with an increased frequency of G>A/C>T transversions, is also in accord with that of GBM^39^. Functionally, many of the GBMO-detected mutations affect genes commonly mutated in GBM, including *IDH2*, *TP53*, and *PTEN*, the chromatin modifiers, *CREBBP*, *SETD2*, *KDM6A*^40^, and *ATRX*, with the latter involved in impaired non-homologous end joining and found in 30% of GBM^41^. Altogether, these data demonstrate that GBMO mimic the mutational landscape commonly seen in GBM (**Fig. 3e,f and Supplementary Table 5**).

**Figure 3.**
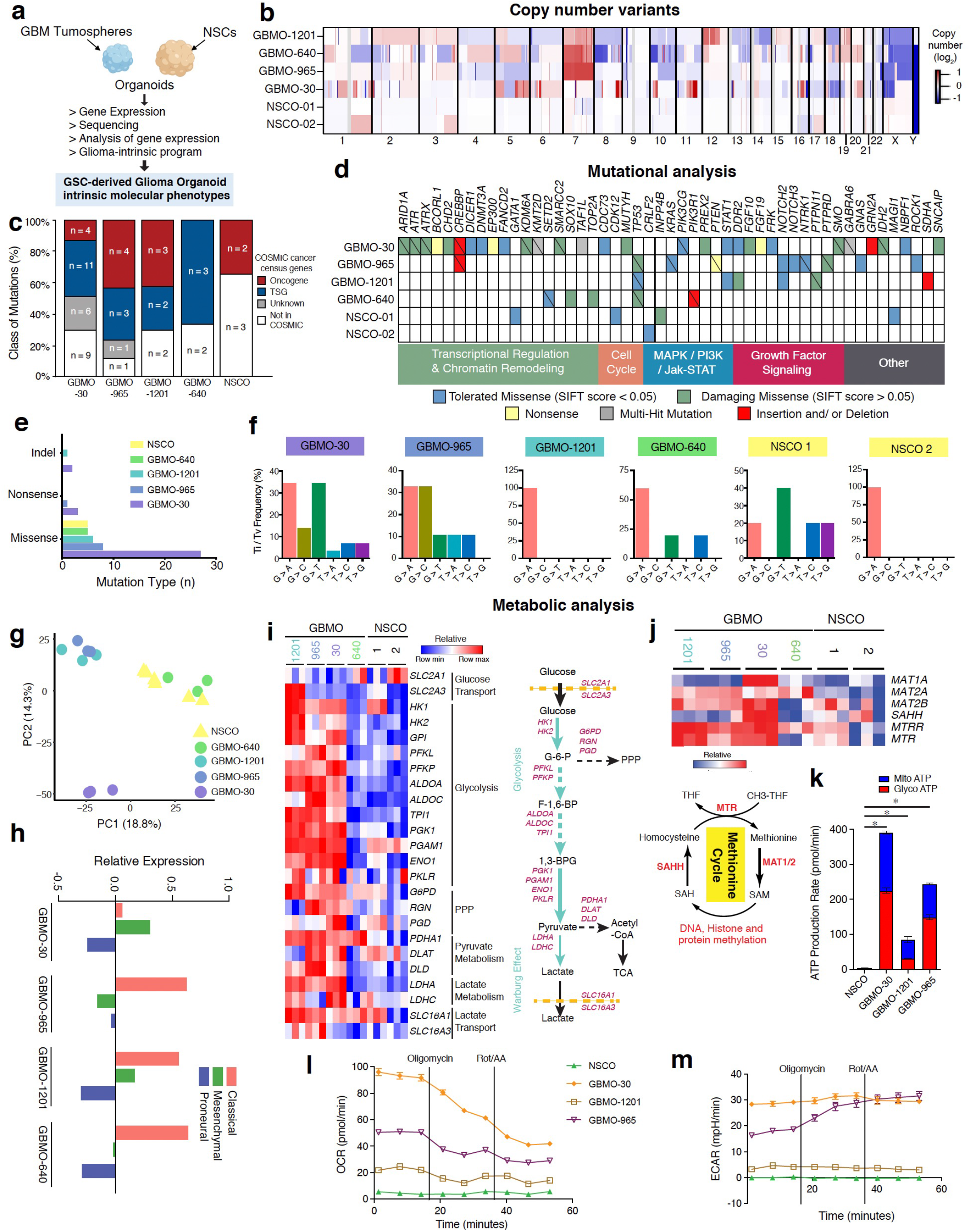
Mutational, transcriptomic and functional metabolic landscape of patient derived GBMO. **(a)** Diagram of genomic sequencing and microarray analysis for GSC-derived glioblastoma organoid intrinsic molecular phenotypes. **(b)** Large-scale copy number variant (CNV) analysis of GBMO-1201, - 640, −965, −30, and NSCO (−01, −02 are technical replicates from different batches). Red, amplifications; blue, deletions. **(c)** Mutational classification in GBMO lines and NSCO, as defined by Catalogue of Somatic Mutations in Cancer (COSMIC). **(d)** Summary of somatic mutation types and functional classification of mutated genes in each GBMO line and NSCO. Complete details are found in **Supplemental Table 3**. **(e)** Quantification of mutation types indicates an enrichment of damaging, coding mutations in GBMO-30, −965, −1201, −640, relative to NSCO. Indel, insertion and/or deletion. **(f)** Transition/transversion (Ti/Tv) proportion comprising single-nucleotide variants for GBMO-30, −965, - 1201, −640, and NSCOs, as identified by TPDS. **(g)** Principal component analysis (PCA) of GBMO and NSCO transcriptomes. **(h)** Relative expression of classifier genes belonging to each GBM molecular subtype **(i)** Metabolomic profiles, as inferred from transcriptomic analysis, of each GBMO line and adjoining summary schematic of classical biochemical pathways. 1,3-BPG, 1,3 Bisphosphoglycerate; F-1,6-BP, Fructose-1-6-Bisphosphate G-6-P, Glucose-6-Phosphate; PPP, Pentose Phosphate Pathway; TCA, Tricarboxylic Acid. **(j)** Expression and schematic of genes involved in the methionine cycle in GBMO and NSCO. GBMO-30, GBMO-965, and GBMO-1201). **(k)** Metabolic analysis with Seahorse XF Real-Time ATP Rate Assay of GBMO-30, 965, 1201 and hNSCO showing mitochondrial ATP (Mito ATP) and glycolytic ATP (Glyco ATP) production rates. Total ATP production: p<0.0001, mito ATP: p<0.0001, glycol ATP: p<0.0001. **(l)** Oxygen Consumption Rate (OCR) and **(m)** Extracellular Acidification Rate (ECAR) were measured among hNSCO and each GBMO.

### Molecular landscapes of GBMO reflect metabolic and molecular alterations found in GBM

Next, we performed gene expression analysis in GBMO and NSCO in which both organoids were maintained in identical culture conditions. Principal component analysis of GBMO global transcriptomes indicated similarity between GBMO-1201 and GBMO-965, as well as between NSCO and GBMO-640, while GBMO-30 clustered by itself, suggesting a distinct expression state from the others (**Fig. 3g**). Using the classifier gene sets^42^, we then calculated subtype scores and found that GBMO-965, −640, and −1201 are most aligned with the classical subtype (**Fig. 3h**). GBMO-30 expressed predominantly mesenchymal subtype genes, in agreement with its absence of SOX2. Consistent with the high metabolic profile of GBM, differential expression analysis indicated that all GBMO except GBO-640 are driven by the Warburg Effect where aerobic glycolysis and lactate fermentation predominate (**Fig. 3i**). Additionally, we found that GBMO showed elevated methionine cycle gene expression, a recently uncovered metabolic pathway essential for glioma stem cells^43^(**Fig. 3j**). Seahorse metabolic analysis confirmed elevated levels of mitochondrial ATP (Mito ATP) and glycolytic ATP (Glyco ATP) production rates in GBMO compared to NSCO (**Fig. 3k-m).**

To better understand GBM gene programs on a global scale, we determined differential expression (DE; defined as log_2_(FC) ± 2 and FDR-adjusted p-value < 0.05) between GBMO and NSCO. We identified 1743 DE genes (DEGs) across GBMO (**Fig. 4a-left and Supplementary Table 6**); unsupervised hierarchal clustering of these DEGs yielded five gene clusters of interest (**gene ontology, Fig.4a-right**). The largest cluster, containing genes almost exclusively upregulated in GBMO, surprisingly was enriched for immune signaling genes associated with immunity, interferon genes (*IFITM1*, *STAT1*, *OAS1*, *IRF9*, *IRF3, IRF1, HLA, NFKB1*), cytokines (*IL6*), as well as hallmark processes of cancer (*IDH1)* and *(MEF/ELF4),* a transcription factor associated with stemness in GBM (**Fig. 4a, blue columns**). The next largest cluster included genes upregulated in GBMO-1201 and −965, and NSCO, suggesting both utilize similar chromatin and Wnt signaling pathways (**Fig. 4a, red columns**). Finally, GBMO-30 was enriched for cell cycle genes, like *ASPM* and *TOP2A*, suggesting it to be the most mitotically active and supporting its highly malignant profile (**Fig 4a, green column**).

**Figure 4.**
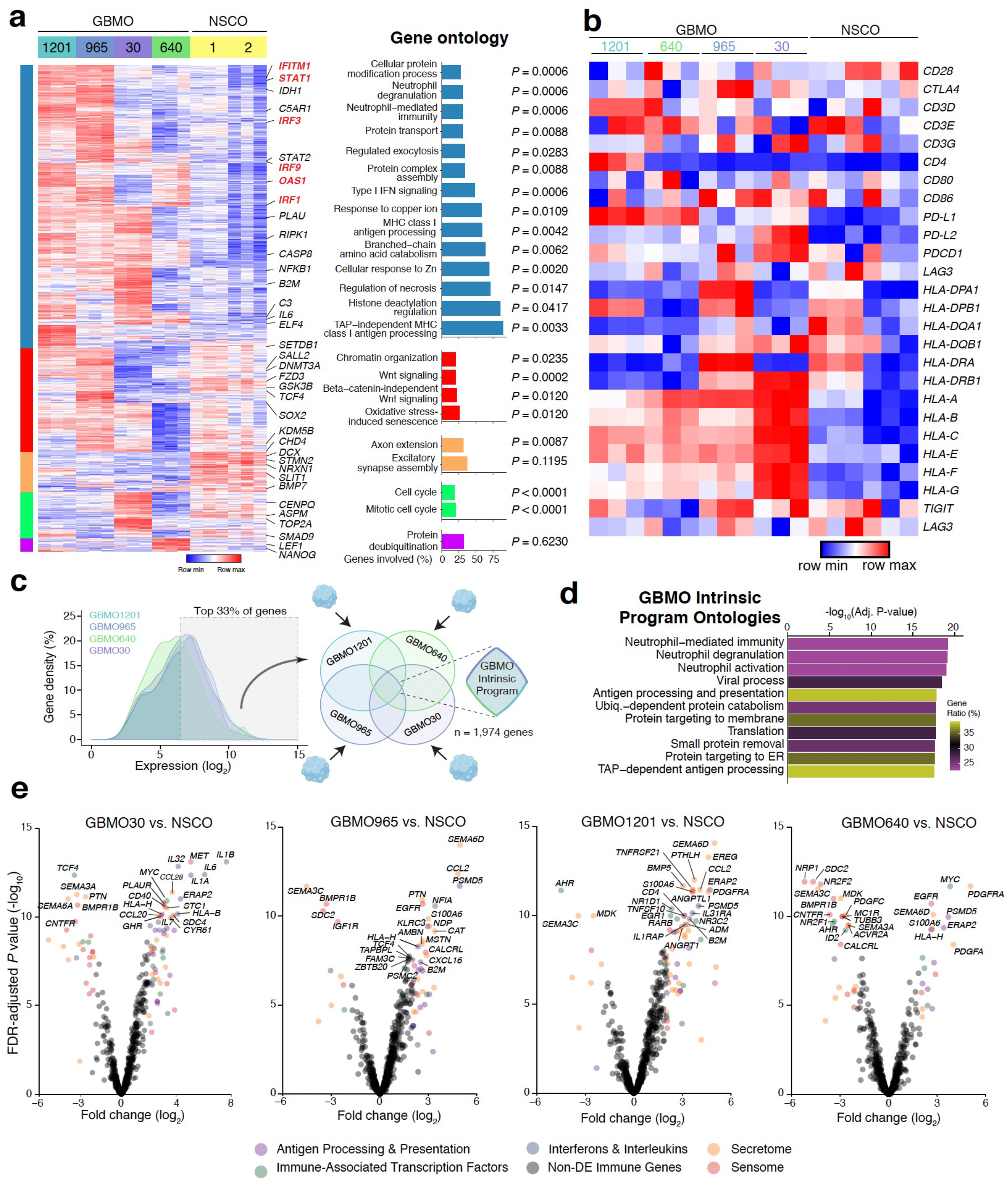
GBMO intrinsic program analysis reveals enrichment of immune genes. **(a)** Hierarchical clustering heatmap of differential gene expression between all GBMO and NSCO with gene ontology for each of the five identified gene clusters. Selected genes from each cluster are highlighted on the right. **(b)** Heatmap for transmembrane proteins of immune system (CDs, PDs, and HLAs) in each GBMO and hNSCO. **(c)** Schematic workflow for identifying genes comprising the GBMO intrinsic program. Briefly, we focused on the shared top 33% of expressed genes from each GBMO line. **(d)** Gene ontology (*right*) enrichment analysis of GBMO intrinsic program genes. **(e)** Volcano plots of differential expression of immune genes for each GBMO line, relative to NSCO. Genes are colored by immune category if differentially expressed (logFC > ±2, adj. *p-*value < 0.05).

In contrast, gene ontology (GO) and pathway analysis of NSCO-specific clusters indicated enrichment for molecules belonging to human neurodevelopment, consistent with the upregulation of known neurogenic genes (*DCX*, *STMN2*, and *NRXN1*) in NSCO (**Fig. 4a, orange column**). We confirmed the diversity of expression of genes proposed to be involved in GBM^25^ which showed heterogenous, but increased expression in all GBMO compared to NSCOs, suggesting that each organoid may have different gene networks that participate in driving their growth (**Supplementary Fig. 4a-b**). We confirmed these transcriptomic results by quantitative polymerase chain reaction (qPCR) in GBMO-30 (**Supplementary Fig. 4c).**

Next, we investigated if the upregulated genes in GBMO are similarly overexpressed in primary gliomas and GBM by leveraging publicly available bulk sequencing data of patient samples from The Cancer Genome Atlas (TCGA) (**Supplementary Fig. 5a**). Evaluation of the topmost upregulated genes in each GBMO by glioma subtype in TCGA revealed a bias toward GBM, opposed to astrocytomas and oligodendrogliomas (17 DE in GBM; 10 DE in Astro; 9 DE in Oligo) (**Supplementary Fig. 5b**). Moreover, examination of these genes by GBM subtype in the TCGA database supported our mesenchymal and classical subtype findings (18 DE in Mes; 19 DE in Cla; 10 in Pro) (**Fig. 3h and Supplementary Fig. 5c**). Since we found distinct regions of proliferation, apoptosis, hypoxia, and quiescent cells in GBMO, we next asked how genes upregulated in GBMOsc reflect gene expression of different GBM histological compartments. To do this, we used the IVY Glioblastoma Atlas Project (Ivy GAP), an RNA sequencing dataset of microdissected GBM structures from 37 patients. While NSCO genes localized to the leading edge, which is known to contain endogenous neural and neoplastic tissue, we observed genes upregulated in GBMOsc to be highly expressed in the leading edge, hyperplastic vessel, and microvascular proliferative domains (**Supplementary Fig.6 and Supplementary Table 7**). Altogether, these results suggest that GSC-derived GBMO reproduce histopathological phenotypes, mutational landscape, expression state, and molecular microanatomy of GBM *in vivo*, providing an opportunity to study the functional relevance of intrinsic molecular pathways at critical stages of GBMO evolution.

### GBMO intrinsic gene expression is enriched for unique glia immune-like molecular programs

One unique advantage of our GBMOsc system is its lack of stromal cells compared to other GBM organoid platforms, which harbor endothelial and immune cells^15^. We leveraged this feature to define an intrinsic glial genetic program, representative of a pre-angiogenic state. To do this, we analyzed the top one-third of expressed genes in each GBMO and focused on the convergence of these genes between all GBMOsc lines to find a shared molecular program. We found 1,974 genes (**Fig. 4c**), which we henceforth refer to as the GBMOsc-intrinsic program (**Fig. 4d**). To gain insights into the functional architecture of this program, we performed pathway and ontology enrichment analyses. The GBMOsc-intrinsic program exhibited enrichment for pathways known to be dysregulated in cancer, such as p53, integrin signaling, glycolysis, and angiogenesis. Gene ontology analysis, on the other hand, highlighted a substantial enrichment for genes involved in immune signaling, consistent with our previous analysis of DEGs (**Fig. 4d and Fig. Supplementary Fig. 7a**). Reasoning that GBM cells may engage in immune interaction with T cells, we next examined the expression of costimulatory/inhibitory pathways^44, 45^. GBMOsc showed increased expression of *PD-L1* in GBMO-640 and GBMO-1201, and *PD-L2* in GBM-30. We also observed elevated expression of MHC class I genes in all GBMOsc, especially GBMOsc-30 (**Fig. 4b**). Together, the expression of classical HLA immune machinery and checkpoint inhibitors by GSC-derived GBMO would, in theory, enable direct interaction with T cells.

Next, we leveraged CIBERSORT^46^, an *in-silico* cell-sorting tool, to determine cell-type immune gene expression across GBMOsc (**Fig. Supplementary Fig. 7b**). GBMO showed similarities in shared immune gene expression, except GBMO-120 that included a sizable proportion of B-cell/plasma cell-related genes, GBMO-640 of M1 macrophage associated genes, and GBMO-30 of mast cell related genes that were different from NSCO, indicating that despite some shared program, the intrinsic immune gene expression is heterogenous and the molecular profile may dictate heterogenus cell-cell interactions with different immune cells. To obtain a more refined view of these immune genes, we evaluated each GBMO compared to NSCO using a collated list of published immune-associated genes (IAGs) from the Immunological Genome Project reference database^47^ and separated these genes into functional clusters representing interleukins/interferons, sensome, secretome, and immune-associated transcription factors (IATFs). This clustering yielded insights into GBMOsc immune-like expression (**Fig. 4e**). For example, most of the upregulated IAGs in GBMO-1201 belong to secretome and interferon/interleukin classes, suggesting it may persistently release immune molecules into the TME, like *CCL2* a chemoattract associated with poor prognosis in GBM. GBMO-640 show expression of *SEMA6D*, *HLA-H* and *ERAP*2, a protease that function by trimming antigenic epitopes for presentation by major histocompatibility complex (MHC) class I molecules. GBMO-30 showed increased *IL6A*, *IL32*, *IL1A*, *ERAP2*, *HLA-B*. Although, there is heterogeneity in immune gene expression, there are some common shared molecules among GBMOsc. To confirm our results and exclude the possibility that our transcriptome data had been confounded by the presence of a few contaminant immune cells, we stained each GBMOsc for immune cell markers such as T cell-specific glycoproteins, CD4 and CD8, and monocyte marker CD68. Indeed, we confirmed their absence in GBMOsc (**Supplementary Fig. 8a**), indicating that these immune-associated genes (IAGs) are intrinsically expressed by GBMOsc cells.

We next sought to correlate our organoids findings in vivo GBM, we turned to two published single-cell RNA-seq data from GBM *in vivo* and examined the expression of key immune genes that we found upregulated. Importantly, the analysis of these genes showed that our GBMO-intrinsic program is observed in tumors *in vivo* and it is not a product of a potential *in vitro* phenomena. We analyzed 25 representative genes of our immune-like genes in these two GBM datasets and found strong in vivo GBM cells intrinsically express immune molecules as shown in our GBMOsc^48^, supporting our observation that GSC-derived GBMOsc recapitulate an immune intrinsic gene expression profile. In vivo, we found GBM genes with immune function can be classified *in vivo* as a) expressed in tumor compartment only (intrinsic) b) expressed in both tumor compartment and infiltrating immune cells; or c) only on immune cells (canonical immune genes), not observed in our GBMOs. These data are consistent with the observation that NSCs and GSC express neuroimmune genes as they interact with immune cells directly, and may modulate the shape the immune contributions by secreting chemokines and cytokines to the TME^20, 22, 23^ ^20, 21^ **(Supplementary Fig. 8b,c).** We correlated the expression of MHC-class I and interferon genes in GBMO and single cell GBM observing that interferon genes like S*TAT1, IRF1, TAP1, IFITM1, IRF3, IRF9* are equally conserved in GBM and our GBMO. To confirm the functional significance of this gene program we examined the effects of IL17 and IFN-γ in the size of GSCs in vitro. We found that IFN-γ but not IL17 decreased the numbers of GBMO-forming GSCs tumorspheres with a concomitant reduction of transcription factor myeloid Elf-1 like factor (MEF), also known as *ELF4*, reduction of *MEF/ELF4* leads to loss of stemness of GBM^49^. **(Supplementary Fig. 9a-c)**. Altogether, these findings provide functional support suggesting that the intrinsic immune program not only can produce molecules able to modulate the TME but also create intrinsic immune vulnerabilities for GBM having intrinsic circuits that can modulate critical genes for stemness in GBM.

### SATB2^+^ and HOPX^+^ progenitor populations drive immune expression in GBM *in vivo*

Having found discrete niches and cell type marker expression in GBMO, we sought to determine which cells were driving the strong expression of immune genes in (**Fig, 4a**). Given that outer radial glia express STAT transcription factors and functional IFN receptors^50, 51^, we hypothesized this population, along with SATB2^+^ cells, which are preferentially born from HOPX^+^ oRG during normal neurogenesis, may contribute to the immune expression of GBMO. To address this, we first analyzed cells that highly expressed SATB2, HOPX, or both together in the dataset and compared the expression of critical ISGs between SATB2^+^, HOPX^+^, SATB2^+^HOPX^+^, and HOPX^-^SATB2^-^ cells (**Fig 5a).** Strikingly, we found elevated levels across several immune genes, including *STAT1*, *IRF3, IRF9*, *IFITM3*, *HLA-A*, and *TAP1* among others (**Fig. 5b**). We repeated this in a separate single-cell GBM dataset and found similar results (**Fig. 5c**). To unbiasedly examine gene signatures of this cell population on a global scale, we performed differential expression between the groups and focused on the most upregulated genes in these cell populations. Pathway enrichment analysis in both datasets revealed these populations shared an enrichment for genes related to epithelial mesenchymal transition, interferon gamma, and interferon alpha signaling (**Fig. 5d, e**). To gain insights to the active immune-related networks of SATB2^+^ and HOPX^+^ cells, we then performed protein-protein interaction analysis, focusing on the topmost upregulated interferon stimulated genes (**Fig. 5f, g**). While these two populations shared some proteins like, *TAP1*, *XAF1,* and *CD74*, there were also some cell-type-specific proteins, including *HLA-C* and *IFITM3* in HOPX^+^ cells and *STAT1* and *STAT2* in SATB2^+^ cells. To confirm these findings from in vivo GBM to our GBMO at the protein shown at RNA level in (**Fig, 4a**), we performed confocal analysis on the expression of IFN-γ receptors and STAT1 in SATB2^+^ and HOPX^+^ cells. Immunostaining revealed that STAT1 and IFNGR1 were indeed expressed by these cells in GBMO-965 and −1201, indicating that our GBMO recapitulate part of the immune gene landscape of HOPX^+^ and SATB2^+^ populations from GBM *in vivo* (**Fig. 5h, i**).

**Figure 5.**
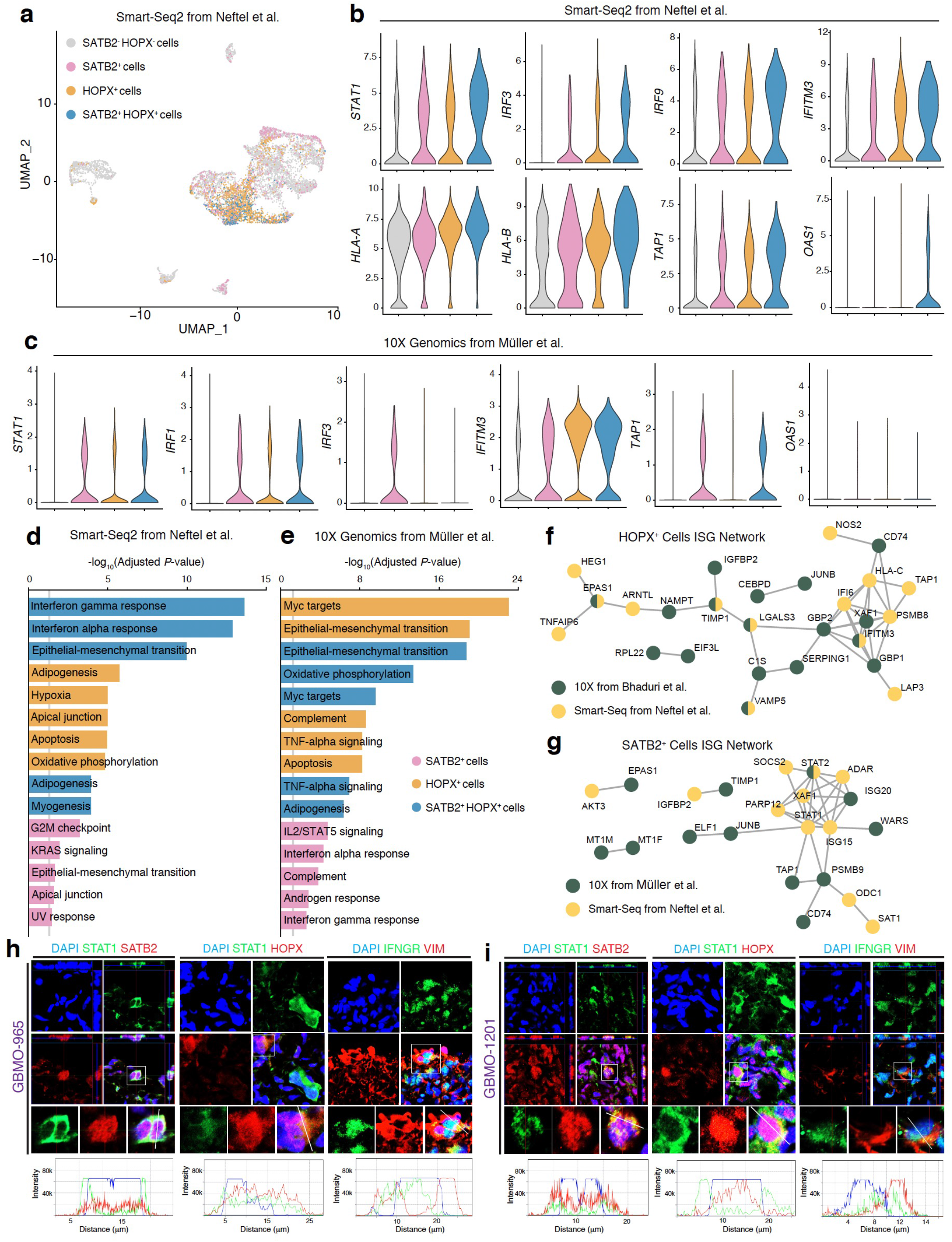
HOPX^+^ and SATB2^+^ cell populations drive immune expression in GBM. **(a)** UMAP plot for *in vivo* GBM single-cell RNA-seq data from Neftel et al^71^. Individual cells are colored according to their high expression (greater than 50% maximum value) of SATB2 (pink), HOPX (orange), both SATB2 and HOPX (blue), or neither (grey). **(b)** Violin plots for representative immune associated genes for each cell population from Neftel et al. **(c)** Violin plots for representative immune associated genes for each cell population from an independent dataset, Muller et al, in which resected GBM were processed for single-cell RNA-seq using 10X Genomics. **(d, e)** Pathway enrichment analysis for genes upregulated in SATB2^+^, HOPX^+^, SATB2^+^HOPX^+^ cells in Neftel et al (d) or Muller et al (e). Grey line denotes 5% level of significance threshold for log-adjusted P-value scores. **(f, g)** Protein-protein interaction network of upregulated interferon stimulated genes in SATB2^+^ and HOPX^+^ cells. Nodes are colored by which dataset they were found to be upregulated. Green, Muller et al; Yellow, Neftel et al. **(h**). Confirmation of expression of interferon related genes on SATB2 and HOPX cancer progenitors. Representative confocal images for immunostaining of STAT1, IFNGR in HOPX and SATB2 populations in 3D reconstruction and z-stacking in GBMO 1201 and 965. Scale bar, 20 μm

## Discussion

In this study, we performed a detailed characterization of the intrinsic properties of patient-derived GBMO from GSCs to significantly expand our understanding of the remarkable ability of GSCs to form a 3D tumor environment from a small, highly malignant group of stem cells (**Supplementary Fig 10**). We demonstrated GSC-derived GBMO recapitulate histological, molecular, metabolic, genetic, and immune features of GBM *in vivo*^52^. Our study importantly shows that GSCs have an encoded developmental memory for 3D tumor evolution that endows them with the ability to give rise to GBMOsc *in vitro*. This is a significant difference from past studies in which GBMO were generated from an established, heterogenous pathological tissue that already has self-organization and a TME. By characterizing the intrinsic programs of GBMO in the absence of immune and vascular contamination, we found an unexpected immune program enriched in GSC-derived cell populations. Accordingly, our GSC-derived GBMO model provides a unique platform to understand basic GBM biology and evolution.

In characterizing the spatial organization of GBMO, we discovered the *in vitro* recapitulation of oRG-like cells in GBMO that have been recently identified in GBM in vivo^31^. These oRG-like GBM cells, distinguished by two oRG-specific markers, p-VIM and HOPX, were present in all patient-derived GBMOsc, despite having distinct transcriptomic profiles (**Fig. 2**), suggesting they may be a ubiquitous cell type in GBM, though more work is needed to conclude this definitively. Underscoring their importance is the intriguing hypothesis which speculates that human GBM initiation originates with aberrant reactivation of the oRG genetic program^50^. In our GBMOsc, these oRG-like tumor cells showed migratory phenotypes and localized to the leading-edge during mitosis. It has also been recognized that a subpopulation of oRG may contain an intrinsic mesenchymal population^53^, perhaps explaining their ability to express immune genes in GBM. Our GBMOsc also recapitulated the presence of SATB2^+^ tumor cells that drive GBM pathology. In normal human brain development, oRG preferentially differentiate into SATB2 upper-layer neurons. Thus, it is tempting to speculate that SATB2 expression may simply be a gene activated downstream of the aberrant deployment of an HOPX oRG-like program, rather than its own malignant progenitor. Indeed, we noted a large portion of cells co-expressing these genes in both single cell RNA-seq datasets. Nevertheless, these novel observations demonstrate that patient-derived GSCs have an encoded ability to give rise to highly malignant progenitors and add to growing evidence that GBM growth and invasiveness may rely on malignant hijacking of the oRG-like and SATB2 gene program. This work also highlights the utility of GBMO for studying GBM biology in a dynamic, 3D cellular system. The insights gained from this level of complexity in recapitulating aspects of tumor biology could not have been achieved by culturing GSCs as tumorspheres *in vitro* (**Supplementary Figure 10**).

There are several models of GBMO, each with different derivations and nomenclature, including 1) genetic activation of oncogenes in normal organoids^4, 12^; 2) invasion of normal iPSC-derived brain organoids by GSCs^13, 16^; 3) GBM organoids derived from fresh pieces of GBM, which include stromal vessels and immune cells^14^; and 4) GSC-derived GBM organoids plus EGF/FGF^15^ and our GBMO system. These models have their own unique advantages and limitations for studying the contributions of the distinct cellular compartments in GBM evolution. Although all GBMO models are limited by the absence of vasculature, we suggest that GSC-derived models may be best suited to model the pre-vascular stages of GBM before initial interactions between tumor, immune, and endothelial cells occur that determine the TME and lead to explosive growth, angiogenesis, and invasion. Therefore, our results support the hypothesis that initial GBM cells would harbor a heterogenous intrinsic immune program that influence, via cell-to cell and paracrine interactions the formation of the TME long before massive vascularization, when major classical immune infiltration happens and underscore that heterogeneity of TME may reside in the intrinsic immune make up of the forming GBM stem and differentiated cells as shown in our GBMOs system.

We capitalized on this opportunity to determine novel tumor-autonomous mechanisms and found strong expression of HLA, tapasin, and interferon genes in GBMO, which using single-cell RNA-seq data, we localized to oRG-like HOPX^+^ and SATB2^+^ cells. We demonstrated the functional role of this pathway and showed that IFN-γ, a cytokine produced by CD8^+^ T cells can target GBMO, causing a downregulation of molecular pathways involved in stemness. Our data support the hypothesis that GBM may have an intrinsic vulnerability for CD8^+^ T cells, including molecules like IFN-γ but this is negated by the powerful influence of GBM cells secreting or harboring molecules like PDL1/2 that induce anergy in infiltrating immune cells. In fact, our model is supported by recent data from a GBM clinical trial that suggest that reverting the T cell exhaustion in GBM by anti-PD1 immunotherapy leads to a restoration of IFN-γ producing T cells and improvement in survival benefit due to T cell targeting of the tumor via IFN-γ^54^. Whether such benefit derives from preferential targeting of the highly invasive and immunogenic HOPX^+^ and SATB2^+^ cell populations (**Fig. 6**) warrants future investigation. Our model should be useful to answer such questions, model tumor specific immune inhibitors, and patient immune cells toward dissecting the immune circuitry of the early TME in a preclinical manner.

**Figure 6.**
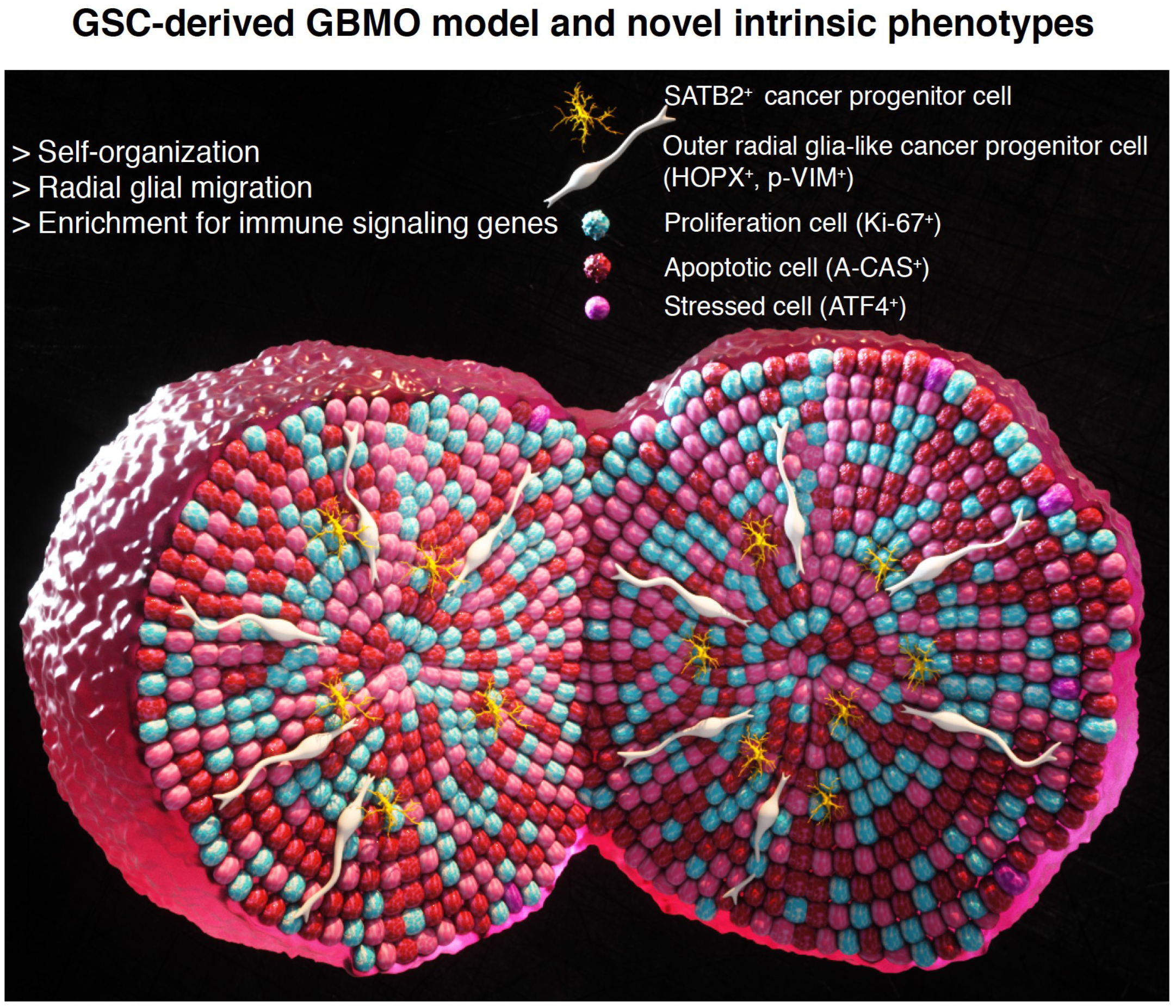
GBM organoid model and novel intrinsic phenotypes. 3D model of a GSC-GBMO organoid summarizing our novel findings, including SATB2 and HOPX oRG-like progenitors that are characterized as bona-fide glioma stem cells. diverse microanatomical domains and hypoxic microenvironments, intrinsic immune program enriched with MHC-class I, interferon and inhibitory pathways in SATB2 and HOPX oRG cells.

Despite the insights into immune biology we found in our GSC-derived GBMO, future GBMO studies will also benefit from incorporating interactions with stromal, vascular, and immune compartments. For instance, the lack of vasculature can be overcome by implantation of organoids orthotopically in mice, though future studies will need to clarify the translational relevance of a xenotransplanted model for studying immune interactions versus co-culturing organoids with immune cells from the same patient^14^. Other future approaches for vascularization could implement microfluidics on a chip with human vascular and immune cells to expand our pre-angiogenic model to a more in vivo-like model. Moreover, whether differences exist in the biology of GBMO seeded from initial, naive GSCs, as modeled here, versus recurrent, therapy resistant GSCs will be another important question to investigate.

In conclusion, we demonstrate GSC-derived GBMO can not only model the unique microanatomy of early 3D tumor formation in the pre-angiogenic/immune stage, but also enable precise identification of intrinsic molecular programs that drive early tumor growth and invasion. To this end, we found a unique neuroimmune and progenitor phenotype that raises questions into the role of the intrinsic immune programs expressed by these malignant progenitors at the earliest stages of TME and tumor evolution. Our GBMO platform thus can serve as a model for future studies seeking to understand both basic GBM biology as well as patient-specific intrinsic vulnerabilities of GSCs in the formation of a complex 3D tumor environment. Much like creating a patient’s tumor avatar, GSC-derived GBMO provide an opportunity for pre-clinical interrogation of patient-derived cells toward personalized treatments for GBM, which currently has a median survival of only 18 months^23^.

## Online Methods

### Generation of organoids and cell culture

Human GBM stem cells (GSCs) were procured from patients and allocated for human research purposes, per the protocols approved by the Institutional Review Boards (IRB) at The Ohio State University Wexner Medical Center and Mayo Clinic^24, 25^. Confirmatory biological properties as GSCs for these lines has been demonstrated elsewhere (**Table 1**). Patient-derived GSCs were isolated and cultured in Neurobasal, 2% B27 supplement without Vitamin A, Glutamax, Antibiotic-Antimycotic (Thermo Fisher Scientific, Waltham, MA, USA), 50 ng/ml human epidermal growth factor (EGF), 50 ng/ml basic fibroblast growth factor (FGF) (R&D Systems, Minneapolis, MN, USA), and heparin (Millipore Sigma, Billerica, MA, USA) in low-attachment cell culture flasks, as previously described^14^. Human neural stem cells (NSC, H9 hESC-Derived, Invitrogen) were cultured on Geltrex in KnockOut DMEM/F12, 2% StemPro Neural supplement, Glutamax, Antibiotic-Antimycotic (Thermo Fisher Scientific), 50 ng/ml human EGF, 50 ng/ml human FGF in low-attachment cell culture flasks, several subclones were generated for the study. For GBM and NSC organoid formation, dissociated single cells were cultured with GBM or NSC growth media for 4-7 days. GBM or normal neurospheres were transferred to Matrigel droplets (BD Bioscience, San Jose, CA, USA) by pipetting into cold Matrigel on a sheet of Parafilm with 3 mm dimples. These droplets were allowed to gel at 37°C and were subsequently removed from the Parafilm. After 4□ days of stationary growth, the tissue droplets were transferred to a spinning bioreactor containing organoid differentiation media including 48% Neurobasal, 48% DMEM/F12, 1% B27 supplement, 0.5% N2 supplement, 1% Glutamax, 0.5% MEM-NEAA, and 1% Antibiotic-Antimycotic^55^. Human iPSCs and iPSC-derived organoids were generated as described previously^55^. All cell lines were handled in accordance with the IBC biosafety practices and relevant ethical guidelines of The Ohio State University College of Medicine, Nationwide Children’s Hospital, and University of Connecticut Health that regulate the use of human cells for research.

### Histology and immunofluorescence

Tissues were fixed in 4% paraformaldehyde for 20 min at 4°C followed by washing in PBS three times for 10 min. Tissues were allowed to sink in 30% sucrose overnight, embedded in OCT compound (Tissue-Tek, Sakura Finetek USA, Torrance, CA, USA), and then cryosectioned at 20 μm. Tissue sections were stained with hematoxylin and eosin, and images were taken with a light microscope (BX41, Olympus, Tokyo, Japan) equipped with a digital camera (DP71, Olympus). For immunofluorescence, sections were incubated successively with 0.25% Triton X-100 and 4% normal horse serum in PBS for 30 min, primary antibodies overnight, and Alexa Fluor 488-, 594-, or 647-conjugated species-specific secondary antibodies for 2 h (Thermo Fisher Scientific). Vectashield Mounting Medium with DAPI (Vector Laboratories, Burlingame, CA, USA) or DAPI (Thermo Fisher Scientific) were used for counterstaining. For single labeling of tissue, the following primary antibodies against the following molecules (immunized species) were used: activated-caspase-3 (559565, BD Pharmingen, San Jose, CA, USA), AHR (ab190797 and ab2769, Abcam), BNIP3 (sc-56167, Santa Cruz Biotechnology), CD4 (550278, BD Pharmingen), CD68 (556059, BD Pharmingen), CD8 (sc-18913, Santa Cruz), CTIP2 (ab18465, Abcam), HOPX (HPA030180, Millipore Sigma), IFNGR1 (MABF753, MilliporeSigma), Ki-67 (ab16667, Abcam), PAX6 (AB_528427, DSHB, Iowa city, IA, USA), phosphorylated-vimentin (D076-3, MBL, Nagoya, Japan), REELIN (MAB5366, Millipore Sigma), SATB2 (ab92446, Abcam), SOX2 (AF2018, R&D systems), STAT1 (610185, BD Bioscience), TBR1 (ab31940, Abcam), TBR2 (ab23345, Abcam), and vimentin (sc7557, Santa Cruz). Images were taken with a confocal laser-scanning microscope (LSM800, Carl Zeiss Microscopy GmbH, Jena, Germany).

### Quantitative analysis of immunostaining

Markers for proliferation and apoptosis (Ki-67 and activated-caspase-3) were quantified and normalized with respect to nuclear DAPI staining, on the outer and inner surface areas of organoid images. Using ImageJ software, quantification of marker staining was performed in 3 equally sized rectangular areas that were overlayed on each outer and inner surface. A minimum of three sections were quantified for each organoid line. The length of relative layer thickness was measured using ImageJ software.

### Targeted parallel sequencing and CNV analysis

A custom capture-based, targeted next-generation sequencing panel, which includes probes covering the coding sequences of 407 cancer-related genes and genome-wide copy number variation (CNV) of backbone targets (Agilent OneSeq 300kb CNV Backbone + custom panel), was utilized in this study. Sequencing libraries were produced using standard methods, barcoded, and sequenced in pools on a HiSeq4000 by the OSU-CCC Solid Tumor Translational Science Shared Resource. Experimental DNA samples were run side-by-side with a human reference DNA sample. Raw sequence reads were aligned with bwa-0.7.13^56^ aln and sample to Homo sapiens genome 1000g v37. Alignment was converted to bam with samtools v1.3.1^57^. After adding read groups, marking duplicates, and sorting with picard 2.4.1^58^, Genome Analysis Toolkit (GATK) v3.6^59^ was used to realign around indels. Mpileup format was generated with samtools v1.3.1 requiring quality > 1 and varscan v2.4.1^60^ ProcessSomatic and Variant Effect Predictor from ensembl^61^ were used to identify tumor-specific variants for each sample. Variants were filtered for location (excluding non-coding variants), coding impact (excluding LOW impact variants – those unlikely to change protein behavior), allele frequency in public databases (excluding common variants found in 1000g^62^ and ExAC^63^), and known variants (excluding variants listed in NCBI dbSNP^64^ but not in COSMIC databases^65^). CNVkit, a python-based copy number calling software^66^, was used to detect the copy number alterations from the samples using a single human “normal” sample as the reference. The log r ratios were calculated for both target and anti-target bins across the genome. Segmentation was performed on the binned data using the circular binary segmentation algorithm. For the heat maps, segments with medians greater than 0.25 were considered duplications and those with medians less than −0.5 were considered deletions.

### Real-Time Quantitative PCR

RNAs were extracted from cell cultures with QIAzol reagent and miRNeasy Mini Kit (QIAGEN, GmbH, Hilden, Germany), following the manufacturer’s protocol. cDNAs were obtained from 500 ng of mRNA using the retrotranscription kit (Thermo Fisher Scientific). Quantitative real-time PCR was performed on 1/20 of the retrotranscription reaction using SYBR Green PCR Master Mix (Thermo Fisher Scientific). Primers were designed to amplify 50- to 200-bp fragments; All primer sequences can be found in Table 2. The qPCR data were assessed using delta-delta CT for evaluating results in the sigmoid region of the amplification curve. For each analysis, samples were normalized by comparison with the housekeeping gene GAPDH. All samples, including “no template” controls, were assayed in triplicate. Each experiment was performed three times with comparable results. Data are expressed as mean ± SEM.

### Transcriptome analysis and differential expression

The GeneChip Human Transcriptome Array 1.0 (also known as Clariom D assays; Affymetrix, Thermo Fisher Scientific Inc.) was used to provide a detailed analysis of the organoid transcriptome. Briefly, 100 ng of total RNA from each of the three samples originally assigned for microarray analysis were used to generate amplified and biotinylated sense-strand cDNA from the entire expressed genome according to the GeneChip WT PLUS Reagent Kit User Manual (P/N 703174, Affymetrix Inc., Santa Clara, CA). cDNA was hybridized to GeneChip Human Transcriptome Array 1.0 for 16 hr in a 45°C incubator, rotated at 60 rpm. After hybridization, the microarrays were washed, and then stained using the Fluidics Station 450 followed by scanning with the Affymetrix GeneChip Scanner 3000 7G, according to manufacturer’s instructions. Raw intensity data was normalized using the quintile normalization of robust multiarray average (RMA) method (performed at the individual probe level). Probes with low-variance were filtered out using the R package genefilter^67^ and annotated to the human genome using the Human Clariom D platform. Transcripts were identified as differentially expressed using the limma package^68^, with a threshold of FDR-adjusted p-value <0.05 and fold change greater than ±2.

### Principal component and subtype analysis, hierarchal clustering, and functional annotation

We reduced the dimensionality of the data by performing principal component analysis (PCA) on the organoid microarray datasets using the prcomp function in R (center = TRUE, scale = FALSE), including only the filtered genes with moderate to high variance. The average relative expression of each set of GBM subtype predictor genes (as defined by TCGA^36^ was quantified from the log2-transformed expression values to determine a relative subtype score for defining each GBMO line. Subtype average expression was normalized by the average log2 expression value of all subtype genes. To identify potential transcriptional modules based on the co-expression of genes in the organoid dataset, unsupervised hierarchical clustering of differentially expressed genes (DEGs) was performed using average linkage and uncentered Pearson correlation on variably expressed genes, as determined by the varFilter() function in limma. Log2-scaled expression values were centered on the median before performing hierarchical clustering. Heatmaps of clustered differential gene expression were then generated. Differentially expressed gene data and the resulting clusters (FC±2, *p*<0.05) were exported for further functional analysis. We made use of Enrichr, which utilizes the Fisher exact test with multiple hypothesis testing correction^69^, to determine gene ontologies, dysregulated pathways, and predicted upstream transcription factors.

### Unbiased search strategy for GSC molecular vulnerabilities

To establish an unbiased GBMO-intrinsic genetic program, normalized intensity values for each GBMO line were first filtered to include only the top one-third of highly expressed genes. The resulting gene lists were compared among GBMO lines and the genes shared by all four lines were the only ones further considered. Gene lists were inputted into Enrichr for enrichment analyses and CIBERSORT^46^ was utilized for *in silico* enumeration of interacting immune cell subsets from GBMO transcriptomic data using default parameters. To further ascertain the immune expression states of each GBMO line, we concentrated on the expression of immune-associated genes, as obtained from ImmPort (http://www.immport.org/immport-open/public/home/home) and InnateDB (http://www.innatedb.com)^70^, using the original DE analyses to ensure all IAGs were encompassed. Plots were produced using ggplot2 package in R. Again, the convergence of the DE IAGs was utilized and then filtered to focus only on known human transcription factors, as specified by http://fiserlab.org/tf2dna_db//index.html. For heatmap generation, data were imported into the online matrix software, Morpheus (https://software.broadinstitute.org/morpheus).

### Processing of single-cell RNA-seq datasets from GBM *in vivo*

Raw read count matrices for Neftel et al^71^ and Muller et al (https://www.biorxiv.org/content/10.1101/377606v1.full72) were downloaded from GSE131928 and UCSC Single Cell Browser (https://cells.ucsc.edu/), respectively. A Seurat object was created for each matrix separately and datasets were scaled. The expression of *HOPX* and *SATB2* were then visualized. In each dataset, an expression cut-off of 50% of maximal gene expression was established and all cells expressing at or above this were deemed to be HOPX^+^ or SATB2^+^. Since we found expression of these genes were not mutually exclusive, we created a third category, HOPX^+^SATB2^+^, for cells expressing both these genes. Cell populations were identified according to these population criteria and the expression of well-known immune associated genes visualized. Upregulated genes in each cell population were then determined for each population using the FindAllMarkers command of Seurat v3^73^. From these upregulated gene lists, pathway analysis was performed using Enrichr. After noting an enrichment of immune-related genes across both datasets, we extracted interferon stimulated genes from the upregulated interferon stimulated genes. We then made use of StringDB (https://string-db.org/) to generate protein-protein interaction networks for the top-25 upregulated ISGs of each cell population and merged these networks from each dataset together.

### Retrospective analysis of gene expression in human gliomas

Gene expression of neurodevelopmental progenitor markers and upregulated GBMO genes were determined across primary patient gliomas and subtypes of GBM tumors, determined through analysis of the National Cancer Institute Repository for Molecular Brain Neoplasia Data (http://betastasis.com/glioma/rembrandt/) and TCGA (https://tcga-data.nci.nih.gov/publications/tcga), respectively. Gene expression localization in structures of primary patient GBM was determined through analysis of Allen Institute of Ivy GAP (http://glioblastoma.alleninstitute.org/). Expression data was downloaded and heatmaps were generated using the matrix visualization software, Morpheus. Heatmaps of Pearson similarity matrices were also generated using Morpheus. Detailed information of heatmaps, including gene names in retained order as in **Supplementary Fig. 7** can be found in **Supplementary Tables 10-14**.

### Metabolic analysis of GBMO with Seahorse technology

For Seahorse Analysis (XFe96, Agilent Technologies), organoids were first dissociated via dissociation reagent. Dissociated cells were washed into warmed Seahorse XF DMEM medium supplemented with 10 mM glucose, 1 mM pyruvate, and 2 mM glutamine and plated at a density of 1×10^5^ cells/well on a poly-L-lysine coated XFe96 Seahorse cell culture microplate. Cells were simultaneously tested for oxygen consumption rate (OCR) and extracellular acidification rate (ECAR) per the manufacturer’s XF Real-Time ATP Rate Assay Kit protocol. Mitochondrial ATP (mitoATP) and glycolytic ATP (glycoATP) production rates were calculated via Agilent Seahorse XF Real-Time ATP Rate Assay Report Generator. ATP Production Rates were analyzed via t-test or ANOVA with Bonferroni posthoc corrections as needed, with significance set at p<0.05.

### Statistical analysis

All data were included for statistical analyses using GraphPad Prism 6.0. Unpaired Student’s t-test (two-tailed) was used for the comparison between two unpaired groups and one-way ANOVA was applied for multi-group data comparisons. The variance was similar between the groups that were being statistically compared. All data met the assumptions of the tests. Survival estimates were calculated using the Kaplan–Meier analysis. Briefly, the expression levels of target genes and patient survival information from the TCGA database were loaded into X-Tile as a tab-delimited text file. By running ‘Kaplan–Meier’ program, the cohort was then divided into two data sets with the optimal cut points generated according to the highest w2-value defined by log-rank test and Kaplan–Meier analyses. Bar graphs were presented as means ± SEM. with statistical significance at *P < 0.05, **P < 0.01, or ***P < 0.001.

## EXTENDED DATA

**Supplementary Figure 1.**
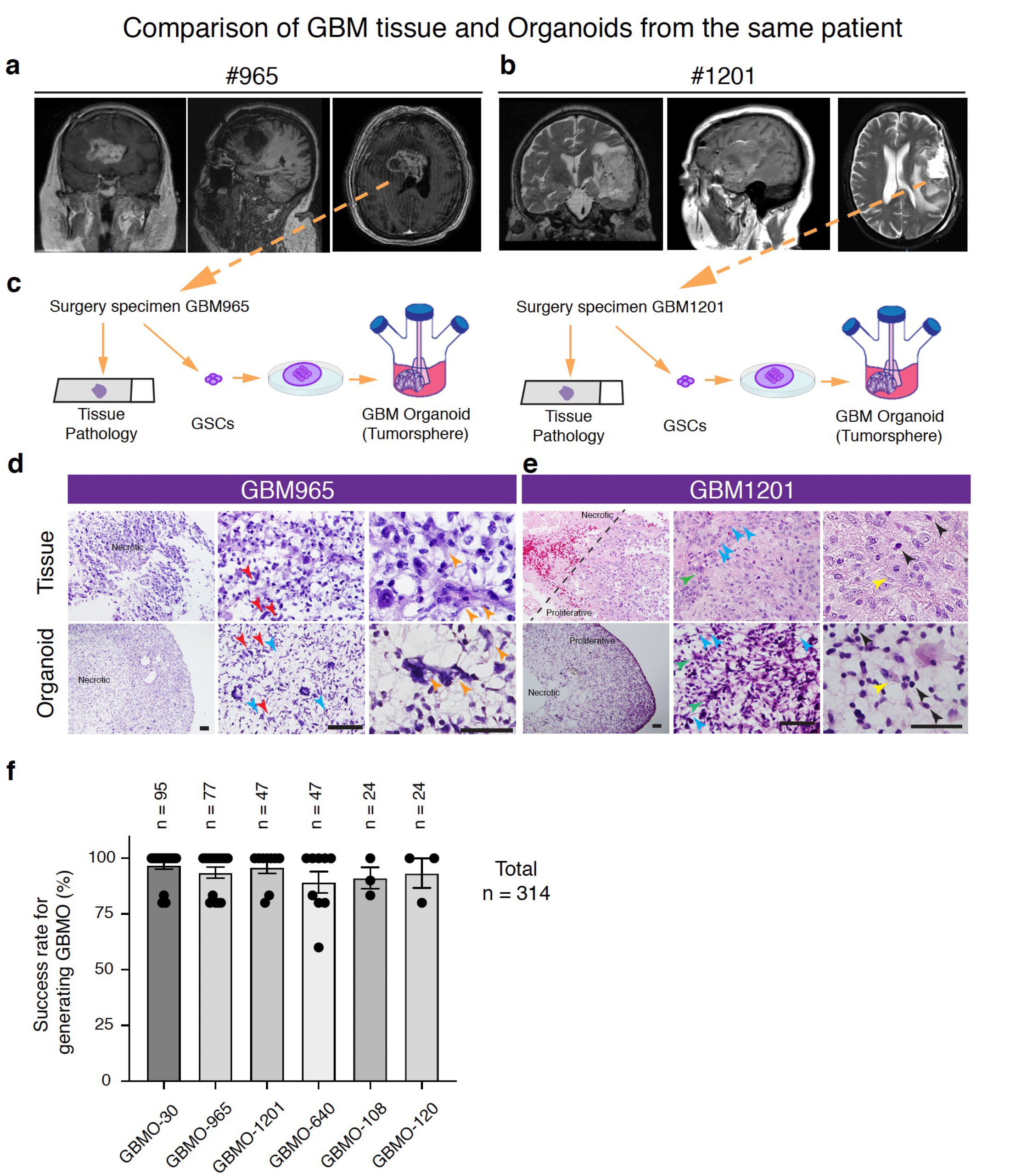
Selected Patient MRI and histopathology of parental tumors and corresponding GBMO (Related to Figures 1 and 2). **(a, b)** Coronal and sagittal MRI slices and tissue pathology of brain tumor patients #965 (left) and #1201 (right). **(c)** Experimental procedure from resected brain tumor to GBM organoids via glioblastoma stem cells (GSCs). **(d, e)** Histological analysis by hematoxylin and eosin (H&E) staining using parental tumor tissue and patient-derived GBM organoid 965 (GBMO-965, left) and 1201(GBMO-1201, right). Red arrows, elongated cells; Orange arrows, syncytia; Blue arrows, fibrils; Green arrows, fusiform cell clusters; Yellow arrows, cells with clear cytoplasm; Black arrows, highly chromatic nuclei. Scale bars, 100 μm. **(f)** Success rate for generating GBMO in different patient derived-glioblastoma stem cells, a total of 226 organoids were generated.

**Supplementary Figure 2.**
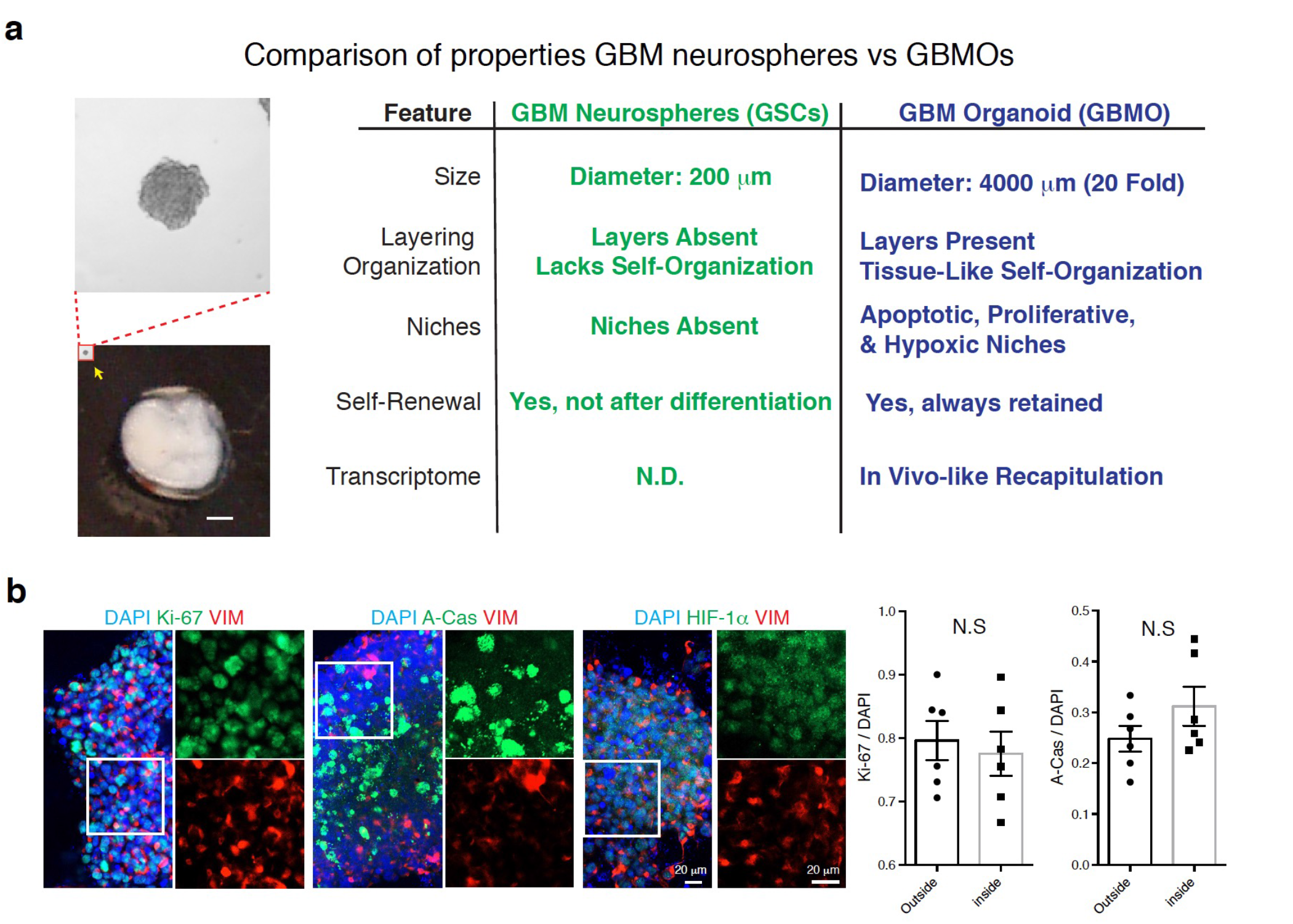
Comparison of in vitro modeling of GBM biology between GBMO and GSC tumorspheres (Related to Figures 1 and 2). **(a)** Comparison of features between GBM tumorsphere and GBM organoid (GBMO). **(b)** Immunocytochemistry for Ki-67, active-caspase-3 (A-Cas), HIF-1α, and vimentin with GBM-30 GSCs on poly-lysine (2D culture). Bar graphs are presented as mean ± SEM. Two-tailed unpaired Student’s *t*-test (N.S. = not significant, *P* > 0.05).

**Supplementary Figure 3.**
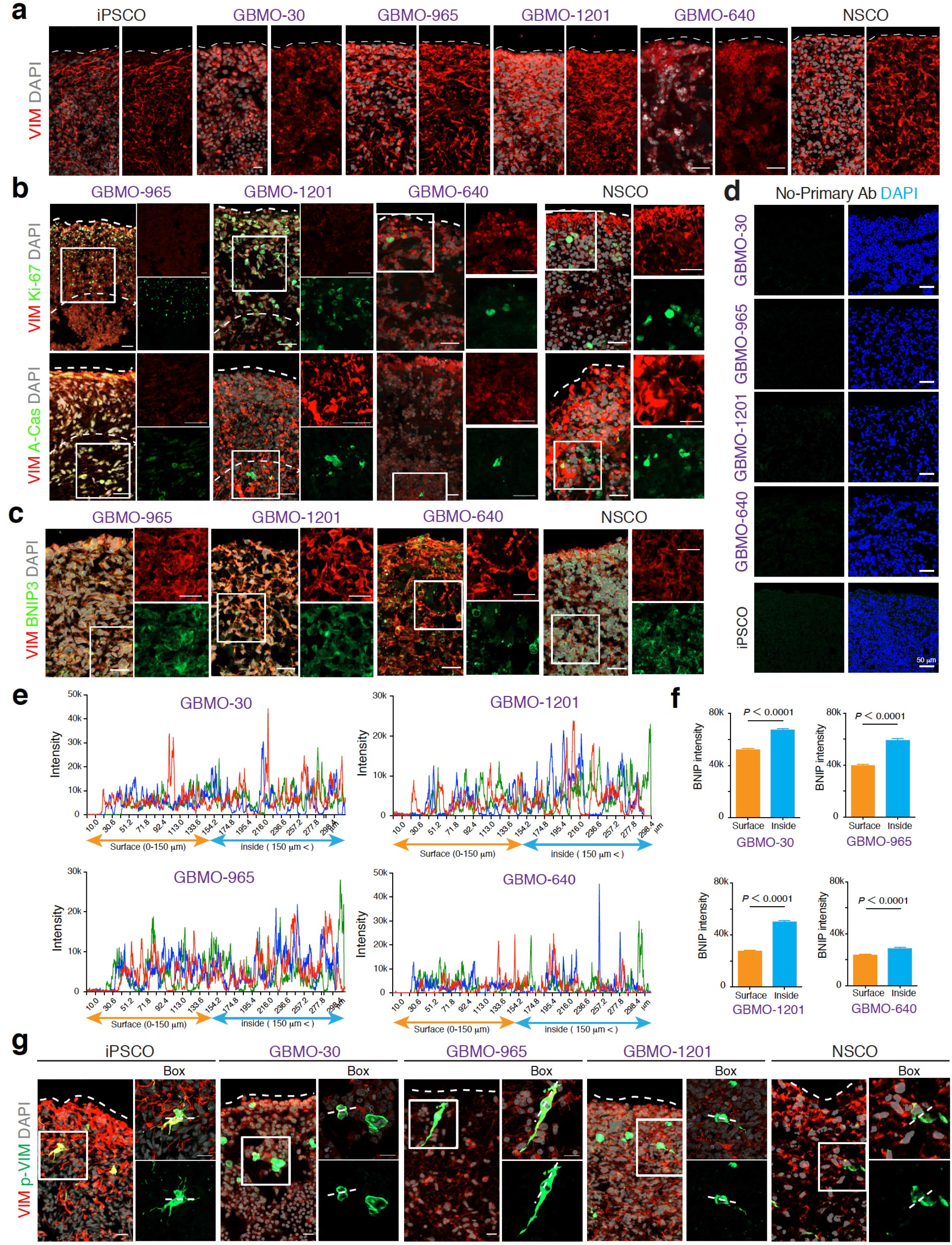
Additional characterization of proliferation, apoptosis, and hypoxia in additional GBMO and hNSCO (Related to Figure 1). **(a)** Representative images of vimentin (VIM) immunostaining in patient-derived organoid from GBM 30, 965, 1201 and 640 (GBMO-30, GBMO-965, GBMO-1201, and GBMO-640) and an organoid derived from human neural stem cells (hNSCO) and human induced pluripotent stem cells (iPSCO). Scale bar, 20 μm. **(b)** Immunohistochemistry for proliferative (Ki-67) and apoptotic (A-Cas) markers in GBMO-965, GBM-1201, GBMO-640, and hNSCO. Scale bar, 50 μm. **(c)** Immunohistochemistry for hypoxia marker (BNIP3) in GBMO-965, GBM-1201, GBMO-640, and hNSCO. Scale bar, 50 μm. **(d)** Negative control (no primary antibody) for each GBMO and iPSCO. **(e)** Hypoxia level (BNIP3 intensity) from surface to inside in GBM-30, −965, −1201, and −640. **(f)** Quantitative analysis for BNIP3 marker intensity between surface (0-150 μm) and inside (>150 μm from surface). **(g)** Sample images of immunostaining for VIM and p-VIM : markers of outer radial glia in iPSCO, GBMO-30, −965, −1201, and neural stem cell organoid (NSCO). Dotted lines highlight cleavage furrow of dividing cells. Scale bar, 20 μm.

**Supplementary Figure 4.**
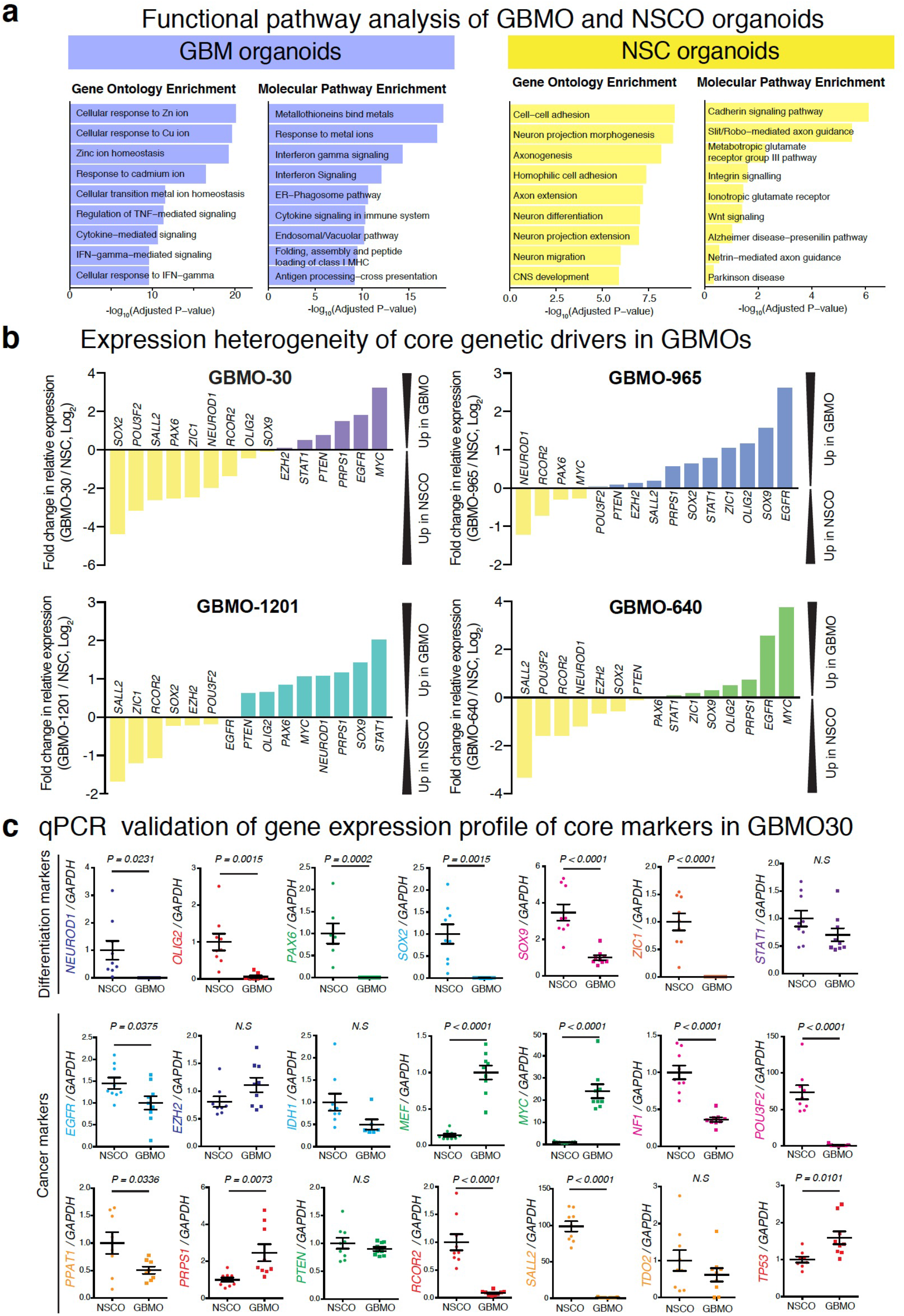
Transcriptomic enrichment for immunity and cancer-related genes in GBMO with confirmatory qPCR in GBMO and NSCO (Related to Figure 2). **(a)** Pathway enrichment using differentially expressed genes from GBMO (left) and NSCO (right). **(b)** Waterfall plot showing expression of notable genes between GBMO-30, −965, −1201, and −640 and NSCO, as detected by transcriptomic analysis. **(c)** Confirmatory qPCR indicates relative expression of differentiation marker genes (top) and cancer-related genes (bottom) between GBMO and NSCO by qPCR. Data are presented as means ± SEM. Two-tailed unpaired Student’s t-test. (*NEUROD1*, *P* = 0.0231; *OLIG2*, *P* = 0.0015; *PAX6*, *P* = 0.0002; *SOX2*, *P* = 0.0015; *SOX9*, *P* < 0.0001; *ZIC1*, *P* < 0.0001; *EGFR*, *P* = 0.0375; *MEF*, *P* < 0.0001; *MYC*, *P* < 0.0001; *NF1*, *P* < 0.0001; *POU3F2*, *P* < 0.0001; *PPAT1*, *P* = 0.0336; *PRPS1*, *P* = 0.0073; *RCOR2*, *P* < 0.0001; *SALL2*, *P* < 0.0001; *TP53*, *P* = 0.0101.). N.S. = not significant, *P* > 0.05.

**Supplementary Figure 5.**
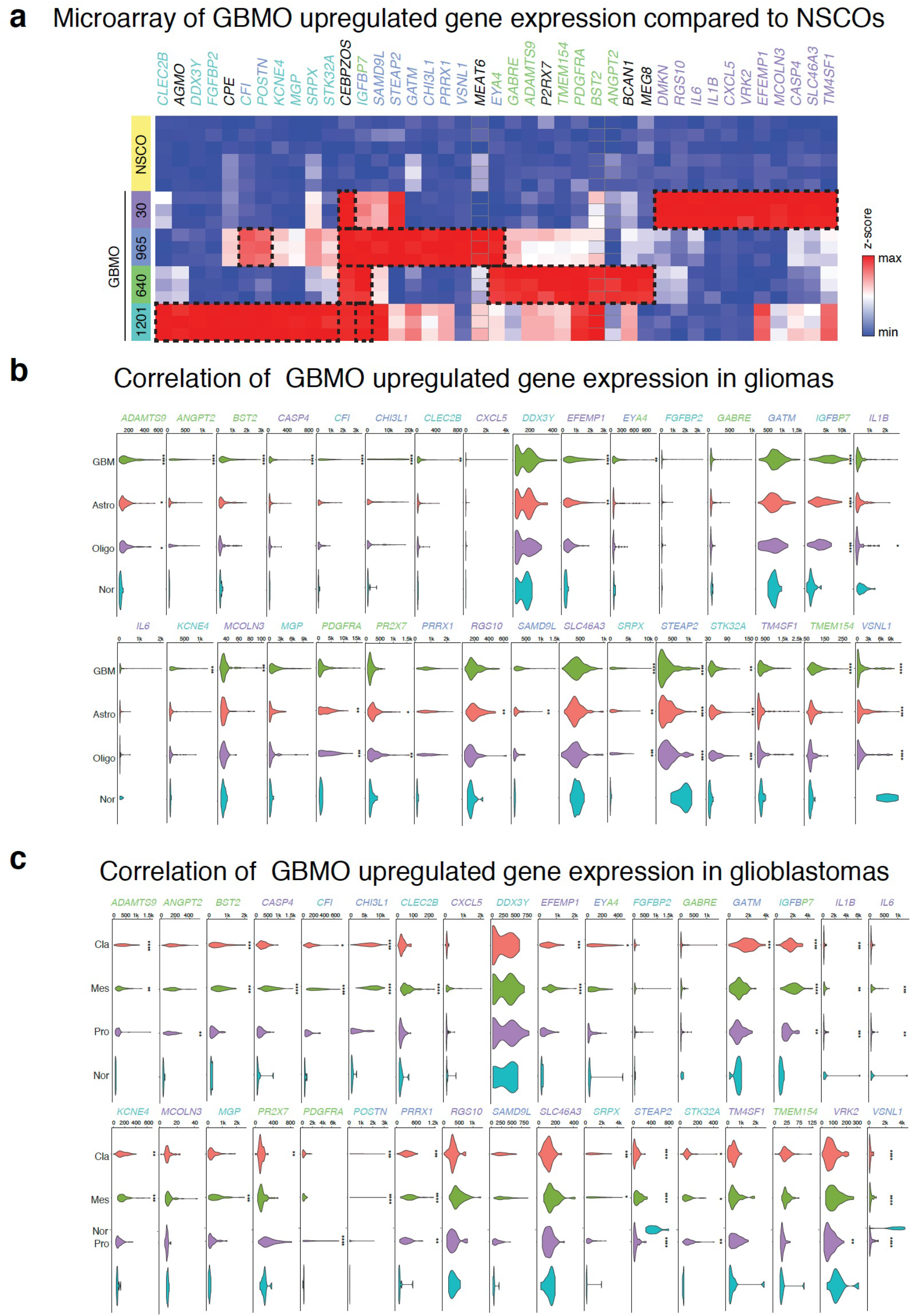
In vivo expression of GBMO-upregulated genes in primary gliomas and GBM subtypes (Related to Figure 2). **(a)** Heatmap of upregulated genes between each GBMO line and NSCO. Dotted boxes denote top twelve genes for each GBMO line. **(b)** Violin plots of upregulated GBMO genes in vitro by glioma subtype in vivo, using publicly available TCGA expression data. Genes colored by which GBMO line they are upregulated in. GBM, Glioblastoma; Astro, Astrocytoma; Oligo, Oligodendroglioma; Nor, Normal. One-way ANOVA followed by Tukey’s test, relative to Normal. (*ADAMTS9*: GBM, *P* < 0.0001; Astro, *P* < 0.05; Olig, *P* < 0.05. *ANGPT2*: GBM, *P* < 0.0001. *BST2*: GBM, *P* < 0.0001. *CASP4*: GBM, *P* < 0.0001. *CFI*: GBM, *P* < 0.0001. *CHI3L1*: GBM, *P* < 0.0001. *CLEC2B*: GBM, *P* < 0.01. *EFEMP1*: GBM, *P* < 0.0001; Astro, *P* < 0.01. *EYA4*: GBM, *P* < 0.01. *IGFBP7*: GBM, *P* < 0.0001; Astro, *P* < 0.0001; Oligo, *P* < 0.0001. *IL1B*: Oligo, *P* < 0.05. *KCNE4*: GBM, *P* < 0.001. *MCOLN3*: GBM, *P* < 0.001. *PDGFRA:* Astro, *P* < 0.01; Oligo, *P* < 0.001. *PR2X7*: Astro, *P* < 0.01; Oligo, *P* < 0.001. *RGS10*: Astro, *P* < 0.01. *SAMD9L:* Astro, *P* < 0.001. *SRPX*: GBM, *P* < 0.0001; Astro, *P* < 0.01; Oligo, *P* < 0.001. *STEAP2*: GBM, *P* < 0.0001; Astro, *P* < 0.0001; Olig, *P* < 0.0001. *STK32A*: GBM, *P* < 0.01; Astro, *P* < 0.001; Oligo, *P* < 0.001. *TMEM154*: GBM, *P* < 0.0001. *VSNL1*: GBM, *P* < 0.0001; Astro, *P* < 0.0001; Oligo, *P* < 0.0001.) **(c)** Violin plots of upregulated GBMO genes by GBM molecular subtype, using publicly available REMBRANDT cohorts. Genes colored by which GBMO line they are upregulated in. Cla, Classical; Mes, Mesenchymal; Pro, Proneural; and Nor, Normal. One-way ANOVA followed by Tukey’s test, relative to Normal. (*ADAMTS9*: Cla, *P* < 0.0001; Mes, *P* < 0.01. *ANGPT2*: Pro, *P* < 0.01. *BST2*: Cla, *P* < 0.001; Mes, *P* < 0.001. *CASP4*: Mes, *P* < 0.0001. *CFI*: Cla, *P* < 0.05; Mes, *P* < 0.0001. *CHI3L1*: Cla, *P* < 0.0001; Mes, *P* < 0.0001. *CLEC2B*: Mes, *P* < 0.0001. *EFEMP1*: Cla, *P* < 0.001; Mes, *P* < 0.0001. *EYA4*: Cla, *P* < 0.05. *GATM*: Cla, *P* < 0.001. *IGFBP7*: Cla, *P* < 0.0001; Mes, *P* < 0.0001; Pro, *P* < 0.01. *IL1B*: Cla, *P* < 0.001; Mes, *P* < 0.01; Pro, *P* < 0.001. *IL6*: Mes, *P* < 0.001; Pro, *P* < 0.01. *KCNE4*: Cla, *P* < 0.01; Mes, *P* < 0.001. *MGP*: Mes, *P* < 0.001. *P2RX7*: Cla, *P* < 0.01. *PDGFRA*: Pro,*P* < 0.0001. *POSTN*: Cla, *P* < 0.001; Mes, *P* < 0.0001. *PRRX1*: Cla, *P* < 0.001; Mes, *P* < 0.0001; Pro, *P* < 0.01. *SRPX*: Cla, *P* < 0.001; Mes, *P* < 0.05. *STEAP2*: Cla, *P* < 0.0001; Mes, *P* < 0.0001; Pro, *P* < 0.0001. *STK32A*: Cla, *P* < 0.05; Mes, *P* < 0.05; Pro, *P* < 0.01. *VRK2*: Pro, *P* < 0.01. *VSNL1*: Cla, *P* < 0.0001; Mes, *P* < 0.0001; Pro, *P* < 0.0001).

**Supplementary Figure 6.**
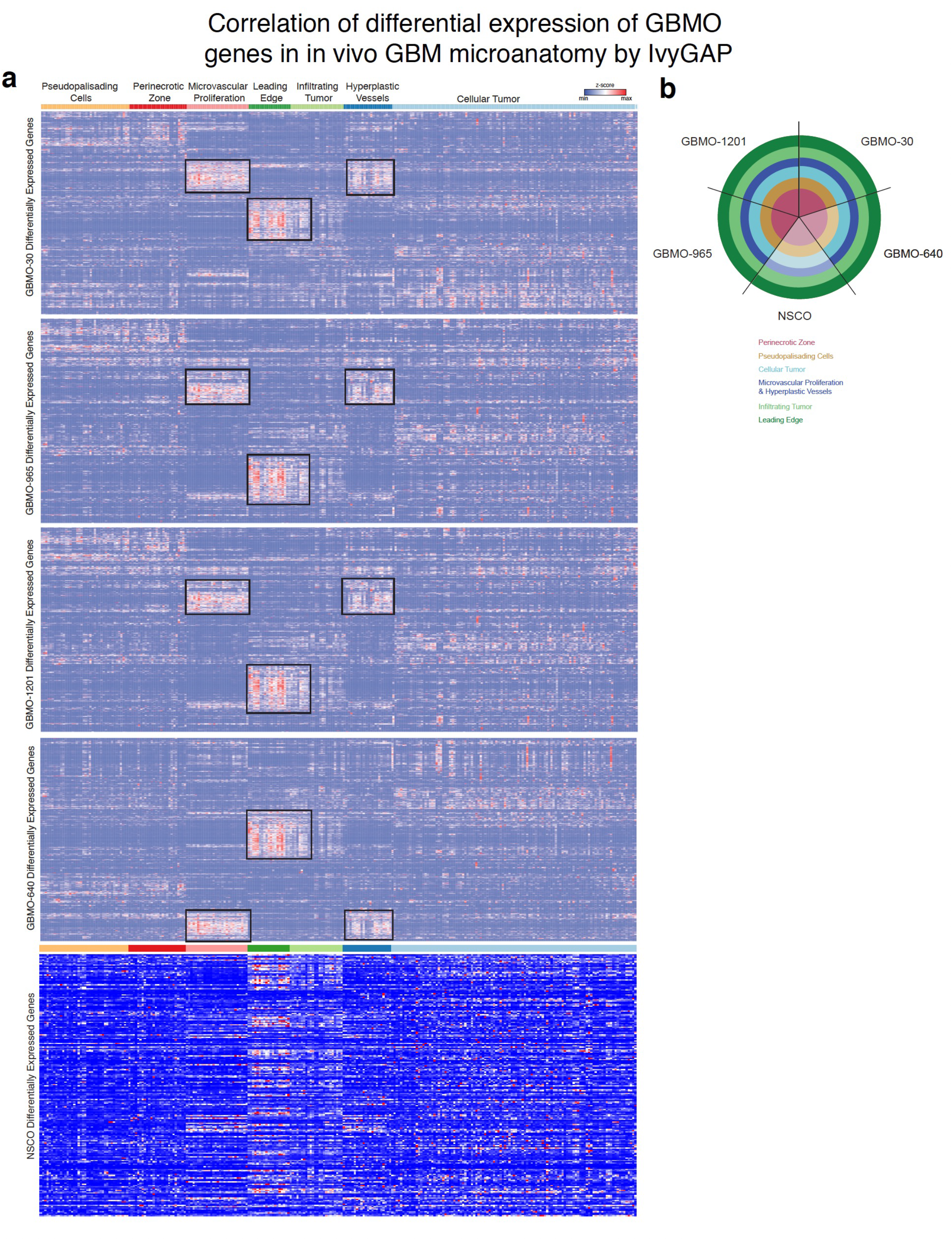
Spatially-resolved heatmap of GBMO-upregulated gene expression in GBM microstructures in vivo (Related to Figure 3). **(a)** Heatmap depicting the spatial expression of upregulated genes in GBMO-30, −965, −1201, and −640 by tumor microanatomy. Complete details of gene names can be found in Supplementary Table 3. RNA-Seq data of tumor structures is derived from IvyGAP. **(b)** Summary figure for these heatmaps. The colors were matched in the circle with the structure colors from the heatmap. Dark colors mean high expression; light colors mean less expression.

**Supplementary Figure 7.**
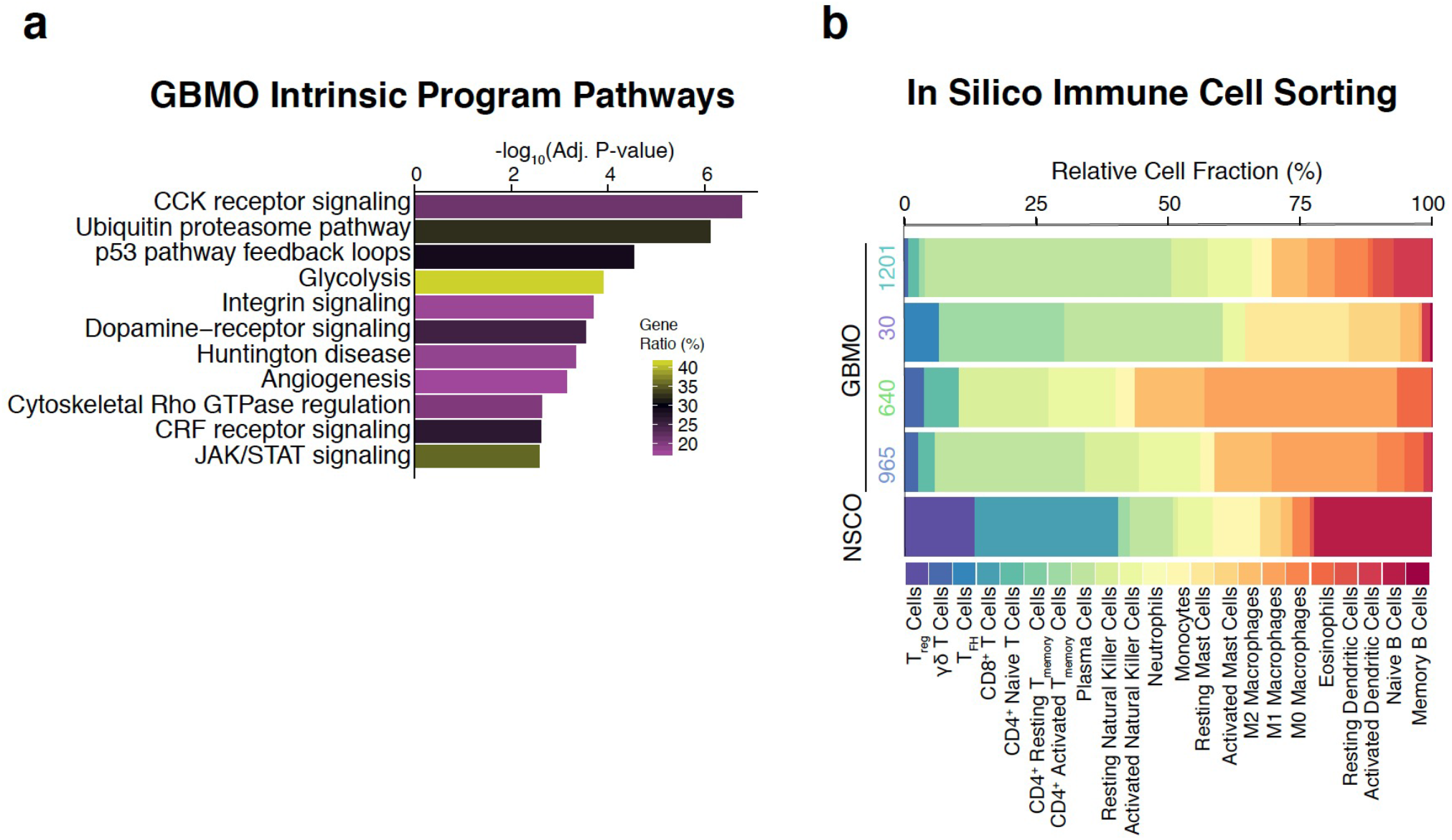
Enrichment of immune genes by GBMO intrinsic program analysis. **(a)** Pathway (*left*) enrichment analysis of GBMO intrinsic program genes. **(b)** *In silico* immune cell transcriptomic analysis for each GBMO line using CIBERSORT to infer immune cell expression signatures within each GBMO line and NSCO.

**Supplementary Figure 8.**
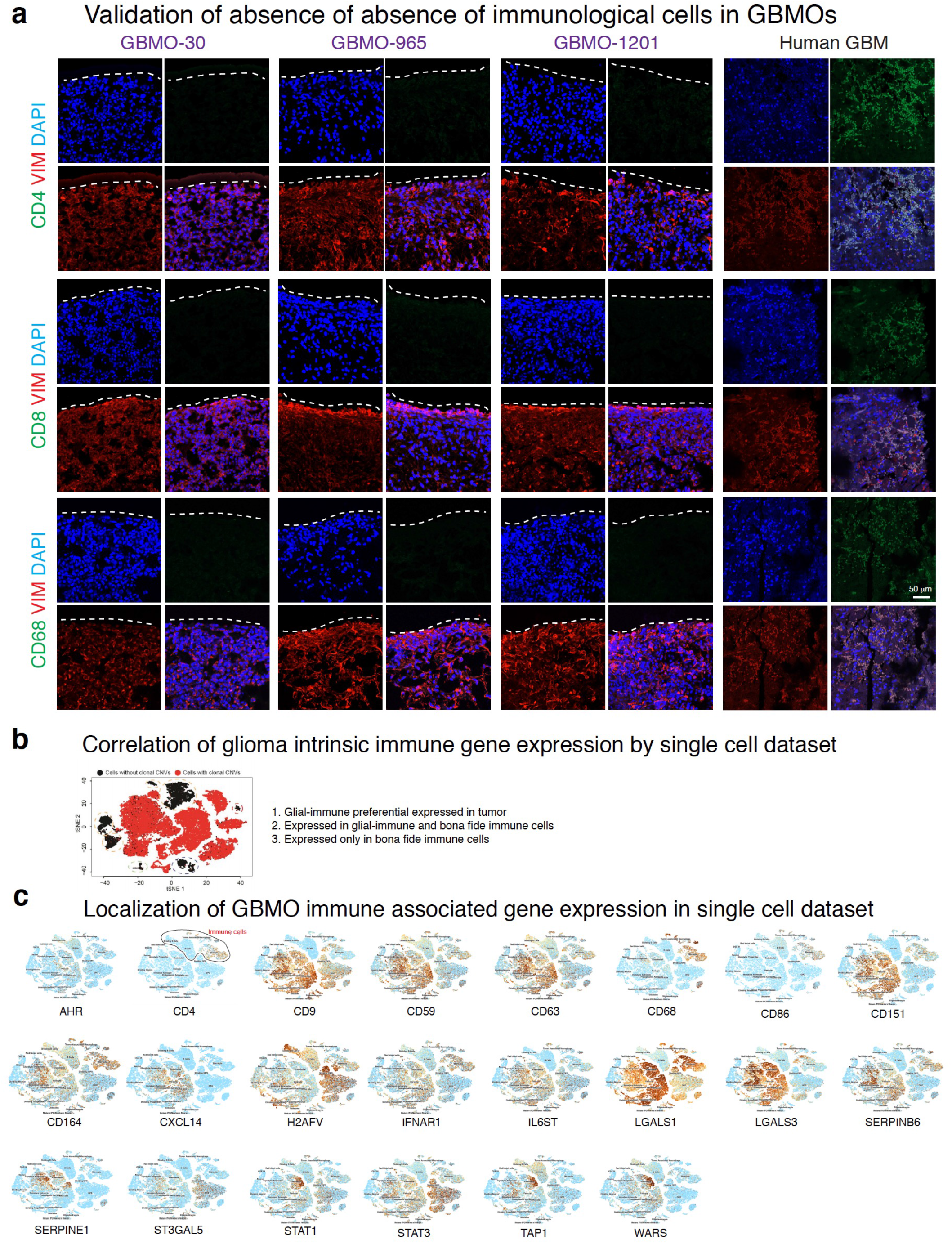
Gene expression for immune system molecules from GBM single-cell RNA-seq and immunostaining for immune cell markers GBMO-30, GBMO-965, and GBMO-1201 and resected GBM tissue (Related to Figures 5). **(a)** Representative confocal images for immunostaining of CD4 (top), CD8 (middle), and CD68 (bottom) in each GBMO line and primary human GBM (positive control). Scale bar, 50 μm. **(b)** Expression of immune-like molecules in the glioma cell vs. infiltrating T cells or macrophages. **(c)** Gene expression in a GBM single-cell RNA-seq for molecules of immune system in vivo GBM.

**Supplementary Figure 9.**
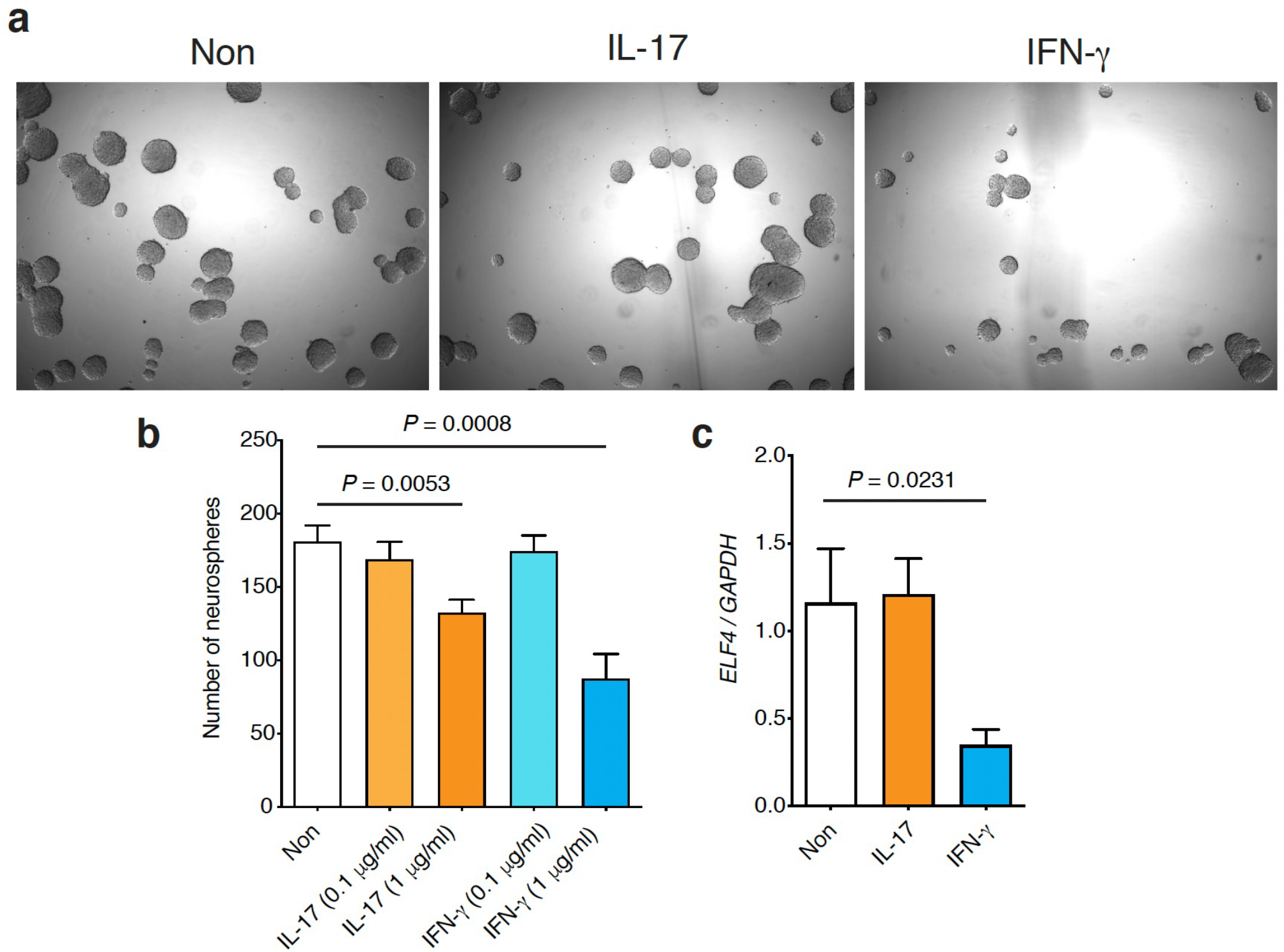
Effects of cytokine in GBMO-forming GSCs self-renewal and *MEF/ELF4* gene expression. **(a)** Neurosphere formation of GBM-30 with IL-17 and IFN-γ. **(b)** Quantification of neurospheres number among IL-17 and IFN-γ -treated GBM-30 GSCs. **(c)** Gene expression of *MEF/ELF4* in GBM-30 with IL-17 and IFN-γ.

**Supplementary Figure 10.**
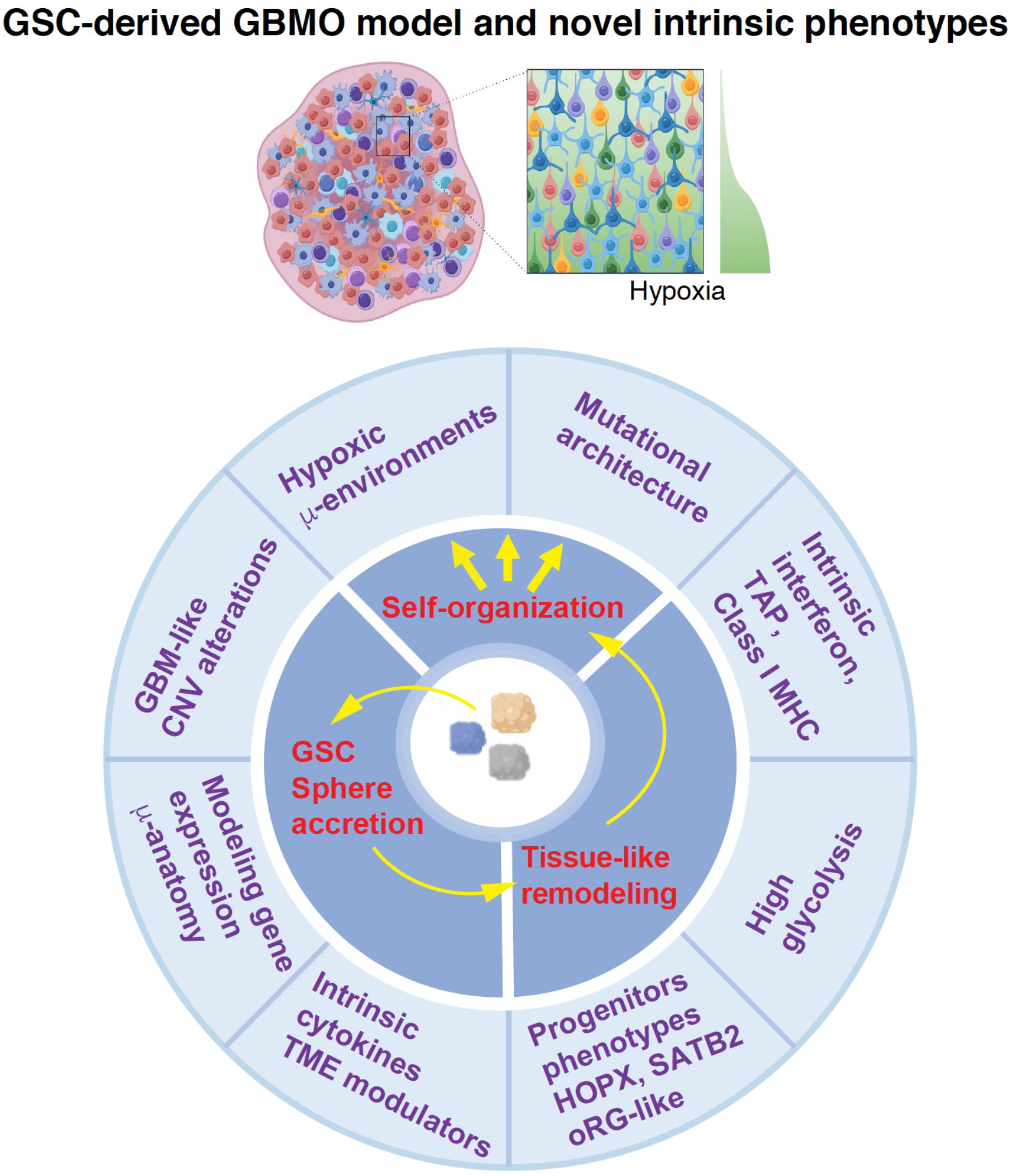
GSC-derived GBMO model and novel intrinsic phenotypes. GSC-GBM organoid model depicting observed novel phenotypes and microanatomy of GBMOs resembling GBMs in vivo.

## Acknowledgments

We kindly thank Sarah Warner for technical expertise in microarrays, Drs. Erica Bell, Jessica Fleming, and Arnab Chakravarti for guidance and design of the targeted next-generation sequencing panel, and The Ohio State Comprehensive Cancer Center Solid Tumor Translational Science Core (Drs. Cynthia Timmers and Tom Liu) and Biostatistics Core (Dr. Amy Webb and Asha Jacob Jannu) for their technical assistance. To Drs. Timothy P. Cripe MD, PhD, and Sofia B. Lizarraga, PhD for their critical reading of the manuscript. The work was supported in part by the NCI (CA016058) and P30 CA016058. AQH is supported by the NIH (CA200399, CA183827, CA195503, CA216855, NS070024), as well as the Mayo Clinician Investigator Award and the State of Florida. VP is supported by grant K24CA 160777. JI was supported by the NRI research Award and The Biogen-UConn Collaboration grant.

## References

1. Guerrero-Cazares, H., Attenello, F.J., Noiman, L. & Quinones-Hinojosa, A. Stem cells in gliomas. Handb Clin Neurol 104, 63–73 (2012).

2. Guerrero-Cazares, H., Chaichana, K.L. & Quinones-Hinojosa, A. Neurosphere culture and human organotypic model to evaluate brain tumor stem cells. Methods Mol Biol 568, 73–83 (2009).

3. Sachs, N. et al. A Living Biobank of Breast Cancer Organoids Captures Disease Heterogeneity. Cell 172, 373–386 e10 (2018).

4. . Bian, S. et al. Genetically engineered cerebral organoids model brain tumor formation. Nat Methods 15, 631–639 (2018).

5. Dietrich, J., Imitola, J. & Kesari, S. Mechanisms of Disease: the role of stem cells in the biology and treatment of gliomas. Nat Clin Pract Oncol 5, 393–404 (2008).

6. Lee, J.H. et al. Human glioblastoma arises from subventricular zone cells with low-level driver mutations. Nature 560, 243–247 (2018).

7. Lathia, J.D., Mack, S.C., Mulkearns-Hubert, E.E., Valentim, C.L. & Rich, J.N. Cancer stem cells in glioblastoma. Genes Dev 29, 1203–17 (2015).

8. Bezzi, M. et al. Diverse genetic-driven immune landscapes dictate tumor progression through distinct mechanisms. Nat Med (2018).

9. Wang, Q. et al. Tumor Evolution of Glioma-Intrinsic Gene Expression Subtypes Associates with Immunological Changes in the Microenvironment. Cancer Cell 32, 42–56 e6 (2017).

10. Azari, H. et al. Isolation and expansion of human glioblastoma multiforme tumor cells using the neurosphere assay. J Vis Exp, e3633 (2011).

11. Ben-David, U. et al. Patient-derived xenografts undergo mouse-specific tumor evolution. Nat Genet 49, 1567–1575 (2017).

12. Ogawa, J., Pao, G.M., Shokhirev, M.N. & Verma, I.M. Glioblastoma Model Using Human Cerebral Organoids. Cell Rep 23, 1220–1229 (2018).

13. Linkous, A. et al. Modeling Patient-Derived Glioblastoma with Cerebral Organoids. Cell Rep 26, 3203–3211 e5 (2019).

14. Jacob, F. et al. A Patient-Derived Glioblastoma Organoid Model and Biobank Recapitulates Inter- and Intra-tumoral Heterogeneity. Cell 180, 188–204 e22 (2020).

15. Hubert, C.G. et al. A Three-Dimensional Organoid Culture System Derived from Human Glioblastomas Recapitulates the Hypoxic Gradients and Cancer Stem Cell Heterogeneity of Tumors Found In Vivo. Cancer Res 76, 2465–77 (2016).

16. Goranci-Buzhala, G. et al. Rapid and Efficient Invasion Assay of Glioblastoma in Human Brain Organoids. Cell Rep 31, 107738 (2020).

17. Imitola, J. Regenerative neuroimmunology: The impact of immune and neural stem cell interactions for translation in neurodegeneration and repair. J Neuroimmunol 331, 1–3 (2019).

18. Zhu, C. et al. The Tim-3 ligand galectin-9 negatively regulates T helper type 1 immunity. Nat Immunol 6, 1245–52 (2005).

19. Watanabe, F. et al. Generation of Neurosphere-Derived Organoid-Like-Aggregates (NEDAS) from Neural Stem Cells. Curr Protoc 1, e15 (2021).

20. Imitola, J. et al. Neural stem/progenitor cells express costimulatory molecules that are differentially regulated by inflammatory and apoptotic stimuli. Am J Pathol 164, 1615–25 (2004).

21. Imitola, J. et al. Directed migration of neural stem cells to sites of CNS injury by the stromal cell-derived factor 1alpha/CXC chemokine receptor 4 pathway. Proc Natl Acad Sci U S A 101, 18117–22 (2004).

22. Merzaban, J.S. et al. Cell surface glycan engineering of neural stem cells augments neurotropism and improves recovery in a murine model of multiple sclerosis. Glycobiology 25, 1392–409 (2015).

23. Al-Kharboosh, R. et al. Inflammatory Mediators in Glioma Microenvironment Play a Dual Role in Gliomagenesis and Mesenchymal Stem Cell Homing: Implication for Cellular Therapy. Mayo Clin Proc Innov Qual Outcomes 4, 443–459 (2020).

24. Banasavadi-Siddegowda, Y.K. et al. PRMT5-PTEN molecular pathway regulates senescence and self-renewal of primary glioblastoma neurosphere cells. Oncogene 36, 263–274 (2017).

25. Banasavadi-Siddegowda, Y.K. et al. PRMT5 as a druggable target for glioblastoma therapy. Neuro Oncol (2017).

26. Chaichana, K.L. et al. Preservation of glial cytoarchitecture from ex vivo human tumor and non-tumor cerebral cortical explants: A human model to study neurological diseases. J Neurosci Methods 164, 261–70 (2007).

27. Qian, X. et al. Brain-Region-Specific Organoids Using Mini-bioreactors for Modeling ZIKV Exposure. Cell 165, 1238–1254 (2016).

28. Fair, S.R. et al. Electrophysiological Maturation of Cerebral Organoids Correlates with Dynamic Morphological and Cellular Development. Stem Cell Reports 15, 855–868 (2020).

29. Morrow, C.S. et al. Vimentin Coordinates Protein Turnover at the Aggresome during Neural Stem Cell Quiescence Exit. Cell Stem Cell 26, 558–568 e9 (2020).

30. Singh, A. et al. The BH3 only Bcl-2 family member BNIP3 regulates cellular proliferation. PLoS One 13, e0204792 (2018).

31. Bhaduri, A. et al. Outer Radial Glia-like Cancer Stem Cells Contribute to Heterogeneity of Glioblastoma. Cell Stem Cell 26, 48–63 e6 (2020).

32. Wang, R. et al. Adult Human Glioblastomas Harbor Radial Glia-like Cells. Stem Cell Reports 14, 338–350 (2020).

33. Suva, M.L., Riggi, N. & Bernstein, B.E. Epigenetic reprogramming in cancer. Science 339, 1567–70 (2013).

34. Jaime-Ramirez, A.C. et al. Humanized chondroitinase ABC sensitizes glioblastoma cells to temozolomide. J Gene Med 19(2017).

35. Tao, W. et al. SATB2 drives glioblastoma growth by recruiting CBP to promote FOXM1 expression in glioma stem cells. EMBO Mol Med 12, e12291 (2020).

36. Cancer Genome Atlas Research, N. Comprehensive genomic characterization defines human glioblastoma genes and core pathways. Nature 455, 1061–8 (2008).

37. Araujo, L.H. et al. Genomic Characterization of Non-Small-Cell Lung Cancer in African Americans by Targeted Massively Parallel Sequencing. J Clin Oncol 33, 1966–73 (2015).

38. Forbes, S.A. et al. COSMIC: High-Resolution Cancer Genetics Using the Catalogue of Somatic Mutations in Cancer. Curr Protoc Hum Genet 91, 10 11 1–10 11 37 (2016).

39. Kandoth, C. et al. Mutational landscape and significance across 12 major cancer types. Nature 502, 333–339 (2013).

40. Liau, B.B. et al. Adaptive Chromatin Remodeling Drives Glioblastoma Stem Cell Plasticity and Drug Tolerance. Cell Stem Cell 20, 233–246 e7 (2017).

41. Koschmann, C. et al. ATRX loss promotes tumor growth and impairs nonhomologous end joining DNA repair in glioma. Sci Transl Med 8, 328ra28 (2016).

42. Verhaak, R.G. et al. Integrated genomic analysis identifies clinically relevant subtypes of glioblastoma characterized by abnormalities in PDGFRA, IDH1, EGFR, and NF1. Cancer Cell 17, 98–110 (2010).

43. Wang, Z. et al. Methionine is a metabolic dependency of tumor-initiating cells. Nature Medicine 25, 825–837 (2019).

44. Hanahan, D. & Coussens, L.M. Accessories to the crime: functions of cells recruited to the tumor microenvironment. Cancer Cell 21, 309–22 (2012).

45. Nakasone, E.S. et al. Imaging tumor-stroma interactions during chemotherapy reveals contributions of the microenvironment to resistance. Cancer Cell 21, 488–503 (2012).

46. Newman, A.M. et al. Robust enumeration of cell subsets from tissue expression profiles. Nat Methods 12, 453–7 (2015).

47. Heng, T.S., Painter, M.W. & Immunological Genome Project, C. The Immunological Genome Project: networks of gene expression in immune cells. Nat Immunol 9, 1091–4 (2008).

48. Wang, Q. et al. Tumor Evolution of Glioma-Intrinsic Gene Expression Subtypes Associates with Immunological Changes in the Microenvironment. Cancer Cell 33, 152 (2018).

49. Bazzoli, E. et al. MEF promotes stemness in the pathogenesis of gliomas. Cell Stem Cell 11, 836–44 (2012).

50. Pollen, A.A. et al. Molecular Identity of Human Outer Radial Glia during Cortical Development. Cell 163, 55–67 (2015).

51. Pollen, A.A. et al. Establishing Cerebral Organoids as Models of Human-Specific Brain Evolution. Cell 176, 743–756 e17 (2019).

52. Hambardzumyan, D. & Bergers, G. Glioblastoma: Defining Tumor Niches. Trends Cancer 1, 252–265 (2015).

53. Eze, U.C., Bhaduri, A., Haeussler, M., Nowakowski, T.J. & Kriegstein, A.R. Single-cell atlas of early human brain development highlights heterogeneity of human neuroepithelial cells and early radial glia. Nat Neurosci (2021).

54. Cloughesy, T.F. et al. Neoadjuvant anti-PD-1 immunotherapy promotes a survival benefit with intratumoral and systemic immune responses in recurrent glioblastoma. Nat Med 25, 477–486 (2019).

55. Lancaster, M.A. & Knoblich, J.A. Generation of cerebral organoids from human pluripotent stem cells. Nat Protoc 9, 2329–40 (2014).

56. Li, H. & Durbin, R. Fast and accurate short read alignment with Burrows-Wheeler transform. Bioinformatics 25, 1754–60 (2009).

57. Li, H. et al. The Sequence Alignment/Map format and SAMtools. Bioinformatics 25, 2078–9 (2009).

58. Picard. Picard - A set of tools (in Java) for working with next generation sequencing data in the BAM format. http://broadinstitute.github.io/picard. http://broadinstitute.github.io/picard.

59. McKenna, A. et al. The Genome Analysis Toolkit: a MapReduce framework for analyzing next-generation DNA sequencing data. Genome Res 20, 1297–303 (2010).

60. Koboldt, D.C. et al. VarScan 2: somatic mutation and copy number alteration discovery in cancer by exome sequencing. Genome Res 22, 568–76 (2012).

61. McLaren, W. et al. The Ensembl Variant Effect Predictor. Genome Biol 17, 122 (2016).

62. Genomes Project, C. et al. A global reference for human genetic variation. Nature 526, 68–74 (2015).

63. Lek, M. et al. Analysis of protein-coding genetic variation in 60,706 humans. Nature 536, 285–91 (2016).

64. Sherry, S.T. et al. dbSNP: the NCBI database of genetic variation. Nucleic Acids Res 29, 308–11 (2001).

65. Bamford, S. et al. The COSMIC (Catalogue of Somatic Mutations in Cancer) database and website. Br J Cancer 91, 355–8 (2004).

66. Talevich, E., Shain, A.H., Botton, T. & Bastian, B.C. CNVkit: Genome-Wide Copy Number Detection and Visualization from Targeted DNA Sequencing. PLoS Comput Biol 12, e1004873 (2016).

67. Huber, W. et al. Orchestrating high-throughput genomic analysis with Bioconductor. Nat Methods 12, 115–21 (2015).

68. Ritchie, M.E. et al. limma powers differential expression analyses for RNA-sequencing and microarray studies. Nucleic Acids Res 43, e47 (2015).

69. Chen, E.Y. et al. Enrichr: interactive and collaborative HTML5 gene list enrichment analysis tool. BMC Bioinformatics 14, 128 (2013).

70. Breuer, K. et al. InnateDB: systems biology of innate immunity and beyond--recent updates and continuing curation. Nucleic Acids Res 41, D1228–33 (2013).

71. Neftel, C. et al. An Integrative Model of Cellular States, Plasticity, and Genetics for Glioblastoma. Cell 178, 835–849 e21 (2019).

72. Muller, S.M., Elizabeth Di Lullo, Aparna Bhaduri, Beatriz Alvarado, Garima Yagnik, Gary Kohanbash, Manish Aghi, Aaron Diaz. A single-cell atlas of human glioblastoma reveals a single axis of phenotype in tumor-propagating cells. https://www.biorxiv.org/content/10.1101/377606v1.full) (2021).

73. Stuart, T. et al. Comprehensive Integration of Single-Cell Data. Cell 177, 1888–1902 e21 (2019).

